# FANCM restrains structural genome evolution and defines a synthetic lethal dependency in *BRCA1*-deficient breast cancer

**DOI:** 10.64898/2026.09.06.749689

**Authors:** Farah Ramadan, Sameer Bikram Shah, Ujwal Subedi, Zachary D. Stephens, Annapoorna Venkatachalam, Neelima Yadav, Vickky Pandit, Sadhana Dubey, Sarah Latka, Hugo Villanueva, Scott H. Kaufmann, Arvind Panday

**Affiliations:** Department of Biochemistry and Molecular Biology, Mayo Clinic, Rochester, MN 55905, USA; Department of Quantitative Health Sciences, Mayo Clinic, Rochester, MN 55905, USA; Department of Oncology, Mayo Clinic, Rochester, MN 55905, USA; Baylor College of Medicine, Houston, TX 77030, USA

**Keywords:** FANCM, BRCA1, synthetic lethality, breast cancer, genomic instability, structural variants, replication stress, tandem duplication, cancer therapy

## Abstract

*BRCA1*-deficient cancers experience persistent replication stress and structural genome instability yet retain the capacity for sustained proliferation, implying reliance on compensatory genome-maintenance mechanisms. Here, we establish BRCA1–FANCM synthetic lethality in human *BRCA1*-deficient breast cancer and exploit temporally controlled FANCM depletion to capture genome evolution over successive cell divisions before declining cellular fitness becomes limiting. We show that FANCM restrains genome-wide structural variation in *BRCA1*-deficient breast cancer cells under endogenous replication stress. FANCM loss amplifies the characteristic *BRCA1*-associated short tandem duplication (TD) phenotype while permitting larger, including megabase-scale, TDs and diverse rearrangements to emerge. Newly emerged TDs preferentially associate with Pol II-occupied regions, and FANCM depletion increases proximity between the replication machinery and elongating RNAPII in *BRCA1*-mutant breast cancer cells, linking FANCM-mediated genome protection to transcription–replication encounters. *BRCA1*-altered human tumors with low FANCM expression recapitulate key features of this phenotype, while genome–transcriptome integration links newly emerged SVs to configuration-dependent local transcriptional changes. Together, these findings establish FANCM as a replication-stress safeguard coupling survival to restraint of structural genome evolution.

## Introduction

BRCA1-deficient breast cancers exhibit profound genome instability arising from defects in homologous recombination and the processing of replication-associated DNA lesions^1–4^. Triple-negative breast cancers (TNBCs) are particularly enriched for BRCA1 dysfunction and frequently display extensive chromosomal rearrangements and characteristic structural variant (SV) patterns^1,5–7^. Among these, tandem duplications (TDs) constitute a prominent genomic feature of breast tumors^8,9^. Genome-wide analyses have further resolved distinct TD size classes associated with different genetic drivers, with BRCA1 deficiency linked to a short-TD population with a modal span of approximately 11 kb^9^. Long-molecule sequencing has further revealed characteristic duplication-containing rearrangement architectures in BRCA1-deficient human cancers and implicated aberrant replication restart in their formation^10^. Despite this persistent burden of replication-associated DNA damage and structural genome instability, BRCA1-deficient cancer cells remain capable of sustained proliferation, implying reliance on compensatory mechanisms that preserve replication under stress. Defining these mechanisms is critical for understanding how BRCA1-deficient cells survive while their genomes continue to evolve, and for identifying vulnerabilities created by BRCA1 loss.

FANCM is an ATP-dependent DNA translocase and replication-fork remodeling factor with a central role in maintaining genome stability during DNA replication^11–13^. FANCM translocase activity promotes replication-fork stability, and loss of this activity increases stalled-fork accumulation and their degeneration into DNA double-strand breaks^12,14,15^. FANCM also functions as a multidomain scaffold and motor protein that coordinates distinct repair pathways at stalled mammalian replication forks^16–19^. Through these activities, FANCM limits pathological processing of stalled replication intermediates and regulates repair pathway choice following replication stress^12,20^. These functions may become particularly important in BRCA1-deficient cells, where protection of stalled/reversed replication forks is compromised and defective fork stability contributes to genome instability^3,21,22^. We therefore hypothesized that BRCA1 deficiency creates an increased requirement for FANCM at stressed replication forks, such that FANCM both supports cell survival and constrains structural genome evolution in the BRCA1-deficient state.

Our previous mechanistic studies provided a foundation for this hypothesis. The Tus/*Ter* system established a site-specific mammalian replication-fork barrier at which BRCA1 regulates homologous recombination and replication-fork repair^2^. Subsequent studies demonstrated that *Brca1*-mutant cells generate characteristic short TDs at Tus/*Ter*-stalled replication forks, establishing a mechanistic link between replication-fork stalling, BRCA1 loss and TD formation^4^. Using this system in *Brca1*-deficient mouse embryonic stem (mES) cells, we further demonstrated that FANCM suppresses mutagenic repair and TD formation at stalled replication forks, while disruption of FANCM function is synthetically lethal with *Brca1* deficiency^16^. These findings suggested that FANCM and BRCA1 provide complementary barriers to mutagenic processing of stalled forks. However, these studies were performed in *Brca1*-deficient mES cells and interrogated repair at a single engineered Tus/*Ter* replication barrier, leaving it unknown whether FANCM performs a comparable genome-protective function in human *BRCA1*-deficient cancer cells. Critically, these studies could not determine whether FANCM continuously restrains structural genome evolution arising from endogenous replication stress over successive cell divisions.

An important unresolved question is whether FANCM protects BRCA1-deficient cells from transcription-associated replication stress. DNA replication and transcription share the same DNA template, creating transcription–replication conflicts (TRCs) when replication forks encounter actively transcribing RNA polymerase complexes. TRCs are increasingly recognized as endogenous sources of replication stress, DNA damage and genome instability^23–25^. These conflicts can be exacerbated by persistent R-loops—three-stranded nucleic-acid structures containing an RNA:DNA hybrid and displaced single-stranded DNA—which can impede replication-fork progression and promote genome instability^26,27^. This is particularly relevant to BRCA1-deficient cancer cells, in which dysregulated transcription and R-loop accumulation have been shown to exacerbate TRCs and compromise genome integrity, identifying excessive transcription–replication collisions as a vulnerability of the *BRCA1*-mutant state^28,29^. Intriguingly, FANCM is mechanistically positioned to counter this vulnerability: purified FANCM can directly displace co-transcriptional R-loops through ATP-dependent branchpoint translocation, while FANCM deficiency increases R-loop accumulation^11,30^. Recent genome-wide analyses further link transcription–replication collisions to TD formation in human cancers, providing a potential route through which unresolved transcription-associated replication stress can be converted into structural variation^31^. Together, these observations raise the possibility that BRCA1-deficient cells become particularly reliant on FANCM to restrain R-loop-associated transcription–replication conflicts. FANCM loss could therefore exacerbate an intrinsic source of replication stress in the BRCA1-deficient state, potentially linking FANCM dependency to the structural genome instability that accompanies BRCA1 loss. Whether this protective function operates in BRCA1-deficient breast cancer and contributes to suppressing genome-wide structural variation, including TD formation at transcriptionally active regions, remains unknown.

Together, these observations raised a broader question: does FANCM provide an adaptive genome-protective function that enables BRCA1-deficient cancer cells to tolerate endogenous replication stress while restraining ongoing structural genome evolution? Several key aspects of this model remained unresolved. Although BRCA1–FANCM synthetic lethality had been identified in *Brca1*-deficient mES cells^16^, whether this dependency operates in human *BRCA1*-mutant breast cancer and, critically, what happens to the surviving cancer genome during sustained FANCM deficiency were unknown. It was also unclear whether BRCA1 deficiency increases FANCM engagement at nascent replication forks as an adaptive response to replication stress. Although FANCM suppresses TD formation at a single engineered Tus/*Ter* barrier, whether this activity extends genome-wide—and whether FANCM loss amplifies the characteristic BRCA1-associated short-TD phenotype, diversifies TD span or generates additional classes of structural variation—had not been determined. Given the intrinsic susceptibility of *BRCA1*-mutant cells to TRCs and the capacity of FANCM to suppress R-loops, a key mechanistic question was whether FANCM loss increases replication–transcription encounters and whether the resulting structural instability preferentially emerges at transcriptionally active genomic regions. Finally, whether FANCM-dependent SV architectures are reflected in human *BRCA1*-altered breast cancers, and whether structural variants emerging during prolonged FANCM deficiency are associated with local transcriptional consequences distinct from the broader cellular response to chronic replication stress, remained unexplored.

Addressing these questions presents an inherent experimental challenge: the *BRCA1–FANCM* synthetic lethal interaction itself limits longitudinal analysis of genome evolution. Constitutive FANCM loss progressively compromises cellular fitness, whereas acute depletion can establish dependency and reveal immediate replication-associated phenotypes but cannot capture structural variants emerging over successive rounds of endogenous DNA replication. We therefore developed a temporally controlled FANCM-depletion system to capture the interval between FANCM loss and declining cellular fitness, enabling newly emerging SVs to be identified before synthetic lethality becomes limiting. This approach transforms synthetic lethality from an experimental endpoint into a temporal framework for interrogating structural genome evolution during chronic FANCM deficiency. Combining longitudinal genome and transcriptome analyses with measurements of FANCM engagement at nascent DNA and replication–transcription encounters allowed us to connect replication-fork biology directly to structural genome evolution and its transcriptional consequences in BRCA1-deficient cancer.

Here, we identify FANCM as an adaptive replication-fork dependency and a genome-wide suppressor of structural genome evolution in BRCA1-deficient TNBC. We show that BRCA1 deficiency is associated with increased FANCM expression and engagement at nascent replication forks and establish FANCM as a synthetic lethality dependency in human BRCA1-deficient breast cancer. By temporally exploiting the interval before synthetic lethality becomes limiting, we uncover a previously unrecognized role for FANCM in continuously restraining structural genome evolution during endogenous replication stress in breast cancer. Chronic FANCM depletion generates reproducible genome-wide SVs, amplifying the characteristic BRCA1-associated short-TD phenotype while permitting larger, including megabase-scale, TDs and diverse rearrangements to emerge. Newly emerged TDs preferentially associate with Pol II-occupied genomic regions, while FANCM depletion increases proximity between the replication machinery and elongating RNAPII in *BRCA1*-mutant cells, linking FANCM-mediated genome protection to transcription–replication encounters. Importantly, *BRCA1*-altered human breast cancers with low FANCM expression recapitulate key features of the experimental duplication phenotype. Integration of genome and transcriptome profiling further distinguishes broad adaptive transcriptional reprogramming from configuration-dependent local expression changes associated with newly emerged SVs. Together, these findings extend FANCM from a regulator of repair at individual stalled forks to a genome-wide determinant of replication-stress tolerance and structural genome evolution, while defining FANCM as a synthetic lethal vulnerability in *BRCA1*-mutant breast cancer.

## Results

### FANCM is preferentially engaged at replication forks in genomically unstable BRCA1-deficient breast cancer

To establish the genomic context in which FANCM may become important in BRCA1-deficient breast cancer, we first examined the relationship between *BRCA1* mutation and genome-wide chromosomal instability across independent breast cancer cohorts. Genomic instability was quantified using the fraction of genome altered (FGA), representing the proportion of the tumor genome affected by somatic copy-number gains and losses. *BRCA1*-mutant tumors exhibited significantly higher FGA than *BRCA1*-wild-type tumors across all three invasive breast carcinoma cohorts **(Fig. 1A; Supplementary Fig. 1A,B)**, demonstrating that BRCA1 deficiency is consistently associated with an elevated burden of genome-wide chromosomal alterations.

**Figure 1.**
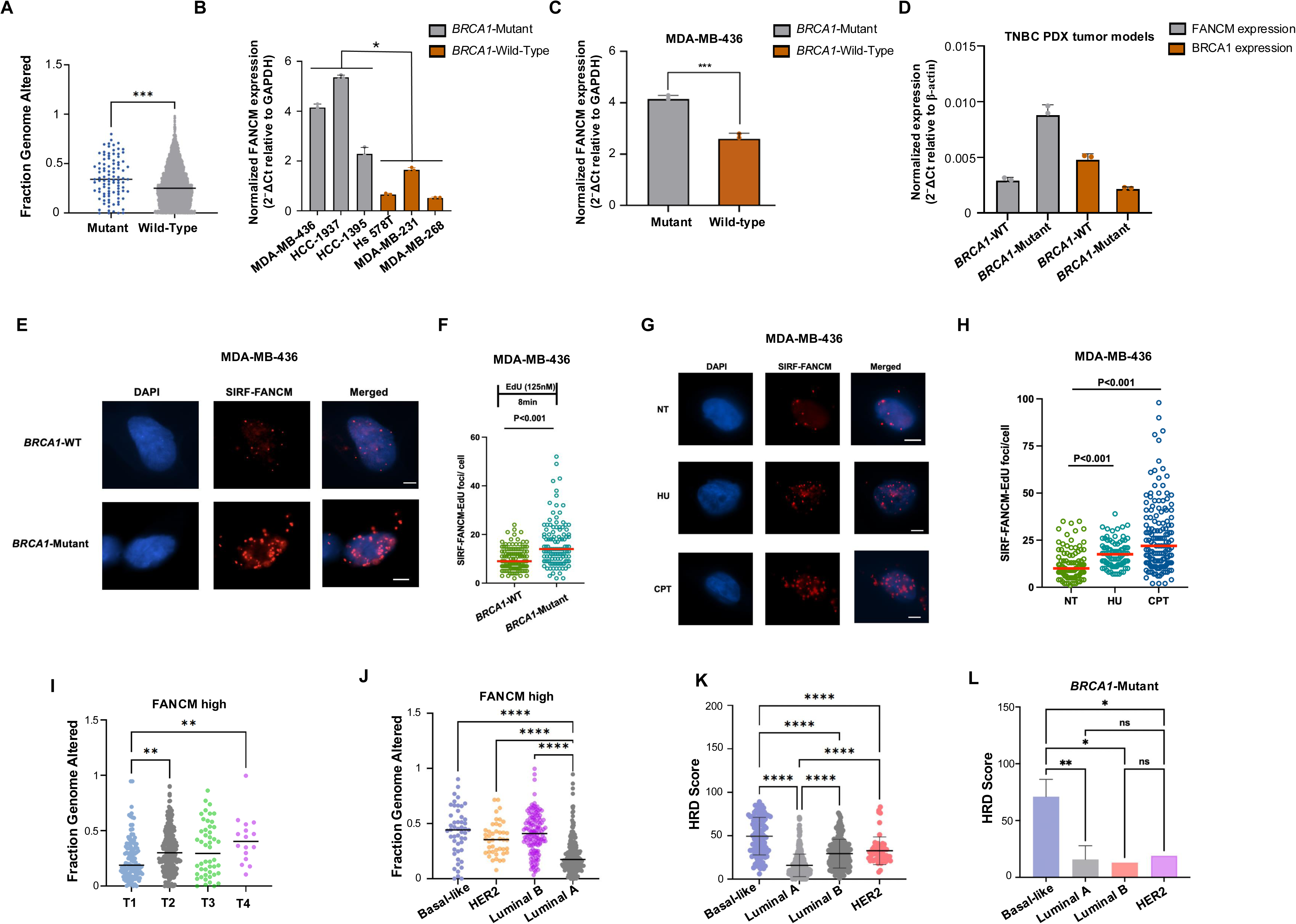
BRCA1 deficiency associates with genomic instability and elevated FANCM expression. **a,** Fraction of genome altered (FGA) levels in *MSK 2025* (n = 3841) cohort relative to *BRCA1* gene status (Wild-type/Mutant). An unpaired Student t-test was used to compare the two cell lines. **b**, Histograms representing FANCM expression levels measured by RT-qPCR in the indicated cell lines, a nested t-test was used to compare the two groups (*BRCA1-*mutant *versus BRCA1-*Wild-type), with three independent cell lines nested within each group. **c**, FANCM expression in parental *BRCA1*-mutant MDA-MB-436 cells and MDA-MB-436 cells complemented with wild-type *BRCA1*. An unpaired Student t-test was used to compare the two cell lines; **b-c,** GAPDH was used for normalization using the 2^−ΔCT^ method. Data are presented as the mean ± SD of 3 independent experiments. **d**, Histograms representing FANCM and BRCA1 expression in *BRCA1*-wild-type and *BRCA1*-mutant TNBC patient-derived xenograft (PDX) tumor models, normalized to β-actin using the 2−ΔCT method. Data are presented as the mean ± SD of 3 independent experiments. Significant differences between the two models were determined using unpaired Student t-test. **e-f** Representative images and quantification of FANCM association with nascent DNA by SIRF (in-situ protein interaction with nascent DNA replication forks) in *BRCA1*-wild-type and *BRCA1*-mutant MDA-MB-436 cells. Cells were pulse-labeled with EdU (125 nM, 8 min); **e**, representative images showing DAPI-stained nuclei (blue), FANCM–EdU SIRF foci (red) and merged channels; **f**, quantification of FANCM–EdU SIRF foci per cell in *BRCA1*-wild-type and *BRCA1*-mutant cells. **g-h** SIRF analysis of FANCM association with nascent DNA following replication stress in *BRCA1*-mutant MDA-MB-436 cells. Cells were pulse-labeled with EdU (125 nM, 8 min) and subsequently left untreated (NT) or treated with hydroxyurea (HU; 0.5 mM) or camptothecin (CPT; 10 µM) for 2 h; **g**, representative images showing DAPI-stained nuclei (blue), FANCM–EdU SIRF foci (red) and merged channels; **h**, quantification of FANCM–EdU SIRF foci per cell under the indicated conditions (scale bars, 5 µm); for **f** and **h**, each point represents an individual cell and red horizontal lines indicate the mean. The *P*-values (two-tailed Mann–Whitney test) are listed at the top. **i-j**, FGA levels across **i**, primary tumor stage (T1-T4), **j**, 4 breast cancer molecular subtypes organized from most to least aggressive: Basal-Like, HER2, Luminal B and Luminal A using TCGA *PanCancer Atlas* dataset for breast invasive carcinoma, irrespective of *BRCA1* mutation status. Only patients having high expression of FANCM (heatmap mRNA expression z-score > 0) were selected. **k-l,** Distribution of Homologous recombination deficiency (HRD) scores across breast cancer molecular subtypes either **k,** irrespective of *BRCA1*-mutation status or **l,** stratified by *BRCA1* mutation status across Basal-like, Luminal A or B, and HER2-enriched tumors. Horizontal lines indicate median values; **i-j,** significance was assessed using one-way analysis of variance (ANOVA), followed by Tukey’s post hoc test for multiple pairwise comparisons and error bars represent standard deviation. For all graphs, asterisks denote significant p-value with ****p<0.0001, ***p<0.001, **p<0.01, *p<0.05, ns = non-significant.

We therefore asked whether FANCM, a replication-stress response factor, is preferentially engaged in the BRCA1-deficient state. FANCM expression was significantly higher across three *BRCA1*-mutant TNBC cell lines (MDA-MB-436, HCC1937 and HCC1395) than in *BRCA1*-wild-type TNBC cells (Hs578T, MDA-MB-231 and MDA-MB-468) **(Fig. 1B)**. This association was recapitulated in an isogenic setting: parental *BRCA1*-mutant MDA-MB-436 cells exhibited significantly higher FANCM expression than MDA-MB-436 cells complemented with wild-type *BRCA1* **(Fig. 1C)**, indicating that FANCM expression tracks with BRCA1 deficiency and is reduced following *BRCA1* restoration. To determine whether this relationship was maintained *in vivo*, we examined FANCM expression in *BRCA1*-mutant and *BRCA1*-wild-type TNBC patient-derived xenograft (PDX) tumor models. FANCM expression was significantly higher in *BRCA1*-mutant PDX tumors, whereas BRCA1 expression was reduced relative to the *BRCA1*-wild-type tumor model **(Fig. 1D)**. Thus, elevated FANCM expression is reproducibly associated with BRCA1 deficiency across genetically distinct TNBC cell lines, an isogenic *BRCA1*-restoration model and PDX tumors.

We next asked whether elevated FANCM expression was accompanied by increased FANCM engagement at ongoing replication forks. FANCM association with nascent DNA was measured using the SIRF (*in situ* protein interaction with nascent DNA replication forks) assay, in which newly synthesized DNA was pulse-labeled with EdU and FANCM proximity to nascent DNA was quantified as FANCM–EdU SIRF foci. Under otherwise unperturbed growth conditions, *BRCA1*-mutant cells exhibited significantly more FANCM–EdU SIRF foci than *BRCA1*-wild-type cells **(Fig. 1E,F)**. This result indicates that the BRCA1-deficient state is associated not only with increased FANCM expression but also with greater localization of FANCM to nascent replication forks, consistent with increased demand for FANCM-mediated fork maintenance under endogenous replication stress.

Because increased FANCM SIRF signal in *BRCA1*-mutant cells could potentially reflect their higher overall FANCM expression, we next asked whether FANCM recruitment to nascent DNA is dynamically responsive to replication stress within the same *BRCA1*-mutant background. MDA-MB-436 cells were pulse-labeled with EdU and subsequently exposed to hydroxyurea (HU) or camptothecin (CPT) to induce replication-fork stalling. HU or CPT treatment significantly increased FANCM–EdU SIRF foci compared with untreated cells **(Fig. 1G,H)**. Thus, FANCM association with nascent DNA increases in response to experimentally induced fork stress independent of comparisons between cell lines with different basal FANCM expression. Together, the SIRF analyses support preferential engagement of FANCM at stressed replication forks and suggest that elevated FANCM activity in BRCA1-deficient cells represents a functional adaptation to their increased replication-stress burden.

We next investigated whether FANCM expression was associated with genomic instability across clinically distinct breast tumor states. Among FANCM-high tumors, FGA increased with tumor progression, with T2 and T4 tumors exhibiting significantly greater genomic alteration than T1 tumors (**Fig. 1I)**. When FANCM-high tumors were stratified by molecular subtype, basal-like tumors displayed significantly greater FGA than Luminal A tumors, while Luminal B tumors also showed elevated genomic alteration relative to Luminal A disease **(Fig. 1J)**. Consistent with this relationship, basal-like tumors exhibited markedly elevated homologous recombination deficiency (HRD) scores compared with Luminal A, Luminal B and HER2-enriched tumors **(Fig. 1K)**. Within the *BRCA1*-mutant setting, basal-like tumors likewise exhibited the highest HRD scores relative to other molecular subtypes **(Fig.1L)**. These analyses position elevated FANCM expression and replication-fork engagement within a tumor context characterized by BRCA1 deficiency, HRD and extensive genomic instability. Collectively, these findings identify FANCM as an adaptive replication-fork response in BRCA1-deficient breast cancer, characterized by both elevated expression and increased engagement with nascent DNA, and provide a functional rationale for testing whether BRCA1-deficient cells become dependent on FANCM for survival.

### BRCA1 deficiency creates a FANCM synthetic lethal dependency that can be temporally interrogated in TNBC cells

The increased expression and replication-fork engagement of FANCM in BRCA1-deficient TNBC cells suggested that FANCM may provide an adaptive response to the elevated replication stress imposed by BRCA1 loss. We therefore asked whether this enhanced FANCM engagement translates into a functional requirement for BRCA1-deficient cell survival. FANCM was depleted using small interfering RNA (siRNA) or short hairpin RNA (shRNA) in three independent *BRCA1*-mutant TNBC cell lines, MDA-MB-436, HCC1395 and HCC1937. Efficient FANCM depletion was confirmed by RT– qPCR **(Supplementary Fig. 2A)**. FANCM depletion markedly reduced colony formation in all three *BRCA1*-mutant cell lines compared with scrambled controls **(Fig. 2A,B)**, demonstrating a conserved requirement for FANCM for long-term survival across independent BRCA1-deficient TNBC models.

**Figure 2.**
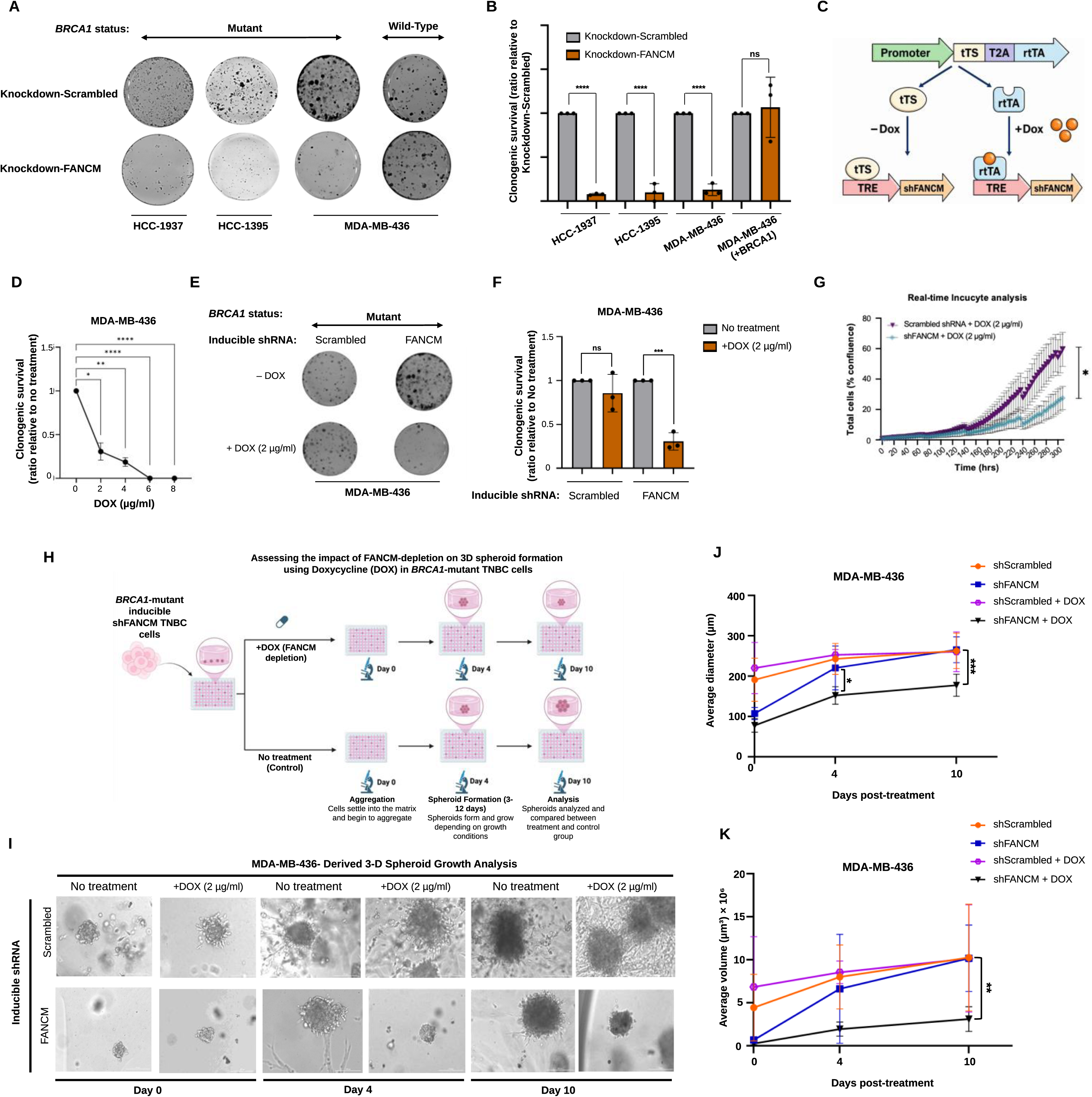
*BRCA1–FANCM* synthetic lethality across 2D and 3D TNBC models and its temporal control. **a-b**, Representative images **(a),** and quantification (**b),** of clonogenic survival assays following FANCM depletion in indicated *BRCA1-*mutant and complemented cells relative to the corresponding scrambled control. Data are presented as the mean ± SD of 3 independent experiments. Asterisks denote significance using unpaired Student t-test (****p<0.0001, ns = non-significant). **c**, Schematic of the doxycycline-inducible shFANCM system. In the absence of DOX, shFANCM expression remains repressed; addition of DOX activates rtTA-dependent transcription enabling temporal FANCM depletion. **d**, DOX dose-response analysis of clonogenic survival in inducible shFANCM MDA-MB-436 cells. Data are presented as the mean ± SD of 3 independent experiments relative to No treatment control. Asterisks denote significant p-value using one-way analysis of variance (ANOVA), followed by Tukey’s post hoc test for multiple pairwise comparisons (****p<0.0001, **p<0.01, *p<0.05). **e-f**, Representative images **(e),** and quantification **(f),** of clonogenic survival assays of inducible scrambled and shFANCM MDA-MB-436 cells cultured with or without DOX (2 µg/ ml) normalized no treatment control. Data are presented as the mean ± SD of 3 independent experiments. Asterisks denote significant p-value using unpaired Student t-test (***p<0.001, ns= non-significant). **g**, Real-time Incucyte analysis of cell confluence during prolonged DOX treatment of inducible scrambled and shFANCM MDA-MB-436 cells. Cell confluence (%) was monitored over time (hours) using live-cell imaging. Data are presented as the mean ± SD of four independent wells per condition. Growth curves were analyzed by Repeated Measures two-way ANOVA using a full factorial model with treatment and time as factors. Multiple comparisons between treatment groups at individual time points were performed using the Holm–Šidák correction. A two-sided P value <0.05 was considered statistically significant. **h-k**, Three-dimensional spheroid assay following inducible FANCM depletion; **h,** Schematic illustration of the experimental workflow; **i,** representative images show spheroid growth at days 0, 4 and 10 (scale bar: 200µm), average spheroid diameters (µm) **(j)**, and average volumes (µm)^3^ **(k)** of spheroids in shScrambled and shFANCM cultures with or without DOX at day 0, 4 and 10. Spheroid diameters were measured using ImageJ (1.54p) and volumes were measured using the formula V=4/3πr^3^, where r stands for radius; data are presented as mean ± SD of 3 independent experiments; asterisks denote significant p-value between shFANCM (with and without DOX) conditions using two-tailed unpaired t-test with Welch’s correction (*p<0.05, ***p<0.001)

To determine whether this dependency was specifically attributable to BRCA1 deficiency, we performed complementation experiments using MDA-MB-436 cells stably expressing wild-type *BRCA1* **(Supplementary Fig. 2A)**. Whereas parental *BRCA1*-mutant MDA-MB-436 cells remained highly sensitive to FANCM depletion, restoration of wild-type *BRCA1* rescued clonogenic survival following FANCM knockdown **(Fig. 2A,B)**. Thus, sensitivity to FANCM loss is determined by the BRCA1-deficient genetic context, establishing conservation of the BRCA1-FANCM synthetic lethal interaction in human TNBC cells.

We next asked what happens to the *BRCA1*-deficient genome when FANCM function is persistently compromised. Although our previous Tus/*Ter* studies established that FANCM suppresses mutagenic repair and tandem duplication formation at a single engineered replication-fork barrier (Tus/*Ter*) in *Brca1*-deficient mouse embryonic stem cells, whether FANCM continuously constrains structural variation across the human cancer genome under endogenous replication stress remained unknown. Addressing this question requires prolonged FANCM depletion over successive cell divisions; however, BRCA1-FANCM synthetic lethality makes conventional long-term depletion inherently limiting.

To overcome this challenge, we developed a doxycycline-inducible shFANCM system in *BRCA1*-mutant MDA-MB-436 cells, creating a temporal window between FANCM depletion and synthetic lethality in which genome evolution could be interrogated **(Fig. 2C)**. Doxycycline significantly reduced FANCM expression in inducible shFANCM cells without significantly affecting FANCM expression in matched scramble controls **(Supplementary Fig. 2B)**. Doxycycline titration produced a dose-dependent reduction in clonogenic survival **(Fig. 2D)**, enabling selection of an induction condition that achieved robust FANCM depletion while retaining sufficient viable cells for prolonged culture and downstream genomic analyses.

The specificity of the inducible system was confirmed using non-targeting scramble and shFANCM cells. Doxycycline exposure had minimal effect on scramble controls but markedly reduced clonogenic survival following inducible FANCM depletion **(Fig. 2E,F)**. Real-time Incucyte analysis further revealed progressive divergence between scramble and FANCM-depleted cultures, with sustained FANCM loss producing a marked reduction in cell confluence at later time points **(Fig. 2G)**. Thus, FANCM depletion causes progressive loss of proliferative fitness rather than immediate cessation of growth, providing an experimentally accessible interval in which the cumulative genomic consequences of FANCM deficiency can be captured.

We further examined this dependency under three-dimensional growth conditions, in which individual cells undergo clonal expansion to form multicellular spheroids. Single MDA-MB-436 cells carrying inducible scramble or FANCM-targeting shRNAs were cultured under spheroid-forming conditions and followed for up to 10 days with or without doxycycline **(Fig. 2H)**. Spheroid outgrowth was quantified longitudinally by measuring both average diameter and average calculated volume. FANCM depletion markedly impaired spheroid expansion, with reductions in both spheroid diameter and volume that became increasingly apparent during prolonged culture **(Fig. 2I,J,K)**. In contrast, doxycycline did not comparably impair spheroid growth in scrambled controls, demonstrating that the growth defect was attributable to FANCM depletion. FANCM is therefore required for *BRCA1*-mutant cell fitness across clonogenic, monolayer and single-cell-derived three-dimensional growth conditions.

Together, these findings establish FANCM as a robust synthetic lethal dependency in BRCA1-deficient TNBC and provide a temporally controlled platform for capturing genome evolution before loss of cellular fitness becomes limiting.

### Chronic FANCM depletion drives structural genome evolution and amplifies BRCA1-associated tandem duplications

To determine whether sustained FANCM loss drives structural genome evolution, we used the inducible system to follow genomic changes during chronic FANCM depletion. Scramble controls and two independently derived inducible shFANCM MDA-MB-436 single-cell clones were maintained under continuous doxycycline exposure for approximately 25 population doublings, allowing replication-associated alterations to accumulate before parallel cultures were harvested for whole-genome sequencing (WGS) and RNA sequencing (RNA-seq) **(Fig. 3A)**

**Figure 3.**
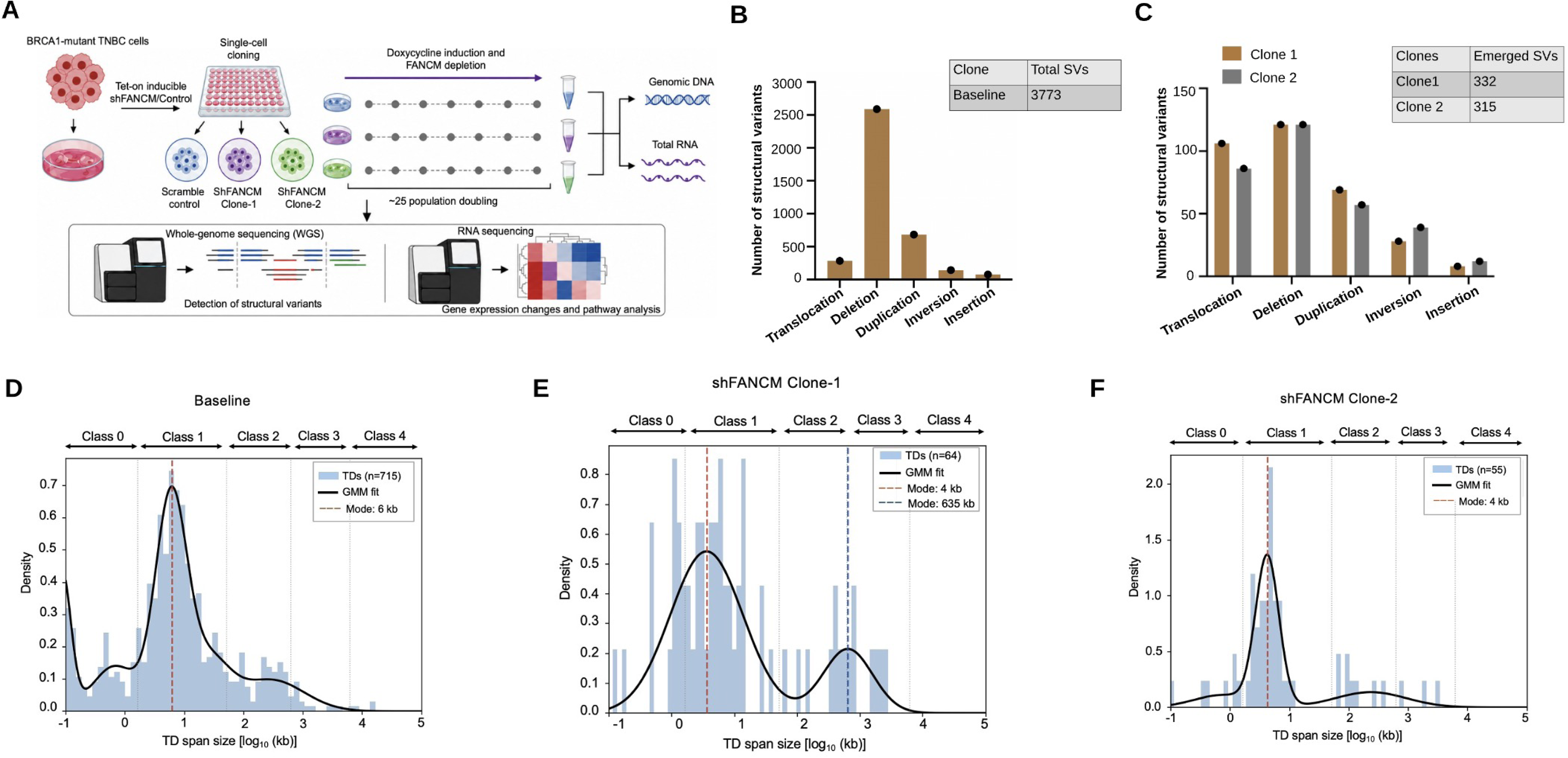
Structural variant and tandem duplication accumulation during chronic FANCM depletion. **a**, Experimental strategy for longitudinal analysis of genome evolution during chronic FANCM depletion. *BRCA1*-mutant MDA-MB-436 cells carrying a doxycycline-inducible scrambled control or two independently derived shFANCM single-cell clones (Clone 1 and Clone 2) were maintained under continuous doxycycline exposure for approximately 25 population doublings. Parallel cultures were harvested for genomic DNA and total RNA and analyzed by whole-genome sequencing (WGS) and RNA sequencing (RNA-seq), respectively. **b**, Baseline structural variant (SV) landscape of *BRCA1*-mutant MDA-MB-436 cells showing translocations, deletions, duplications, inversions and insertions. **c**, SVs newly emerging during chronic FANCM depletion in Clone 1 and Clone 2. **d–f**, TD span-size distributions and Gaussian mixture model (GMM) fits for the baseline *BRCA1*-mutant genome **(d)**, shFANCM Clone 1 **(e)**, and shFANCM Clone 2 **(f)**. TD span is plotted as log10(kb), with GMM-derived size classes (Classes 0–4) indicated above each distribution. Black curves represent GMM fits and colored dashed lines indicate GMM-derived modes.

We first defined the pre-existing SV landscape against which FANCM depletion-induced events could be identified. The baseline *BRCA1*-mutant genome contained 3,773 structural variants, dominated by deletions, with additional translocations, duplications, inversions and insertions **(Fig. 3B)**. Using this baseline as reference, WGS identified 332 and 315 additional SVs in Clone 1 and Clone 2, respectively, following chronic FANCM depletion **(Fig. 3C)**. These events were absent from the baseline dataset and therefore represent SVs that emerged during the FANCM-depletion interval. Deletions and translocations were the most abundant newly emerged classes in both clones, followed by duplications, inversions and insertions. The comparable burden and composition of additional SVs across two independently derived clones demonstrate that chronic FANCM loss reproducibly drives structural genome evolution.

Given our previous observation that FANCM suppresses TD formation at a site-specific Tus/*Ter* replication barrier, we next asked whether chronic FANCM depletion affects the genome-wide TD landscape under endogenous replication stress. The baseline MDA-MB-436 genome contained 715 pre-existing TDs, with Gaussian mixture modeling identifying a dominant population centered at approximately 6 kb **(Fig. 3D)**. Chronic FANCM depletion generated 64 and 55 additional TDs in Clone 1 and Clone 2, respectively.

Newly emerged TDs retained a prominent short-duplication population in both clones. Clone 1 exhibited a principal mode near 4 kb together with a second mode at approximately 635 kb, producing a broadened bimodal distribution **(Fig. 3E)**. Clone 2 similarly retained a dominant mode near 4 kb, accompanied by lower-frequency events extending toward larger genomic spans **(Fig. 3F)**. Thus, FANCM loss does not replace the underlying BRCA1-associated TD phenotype but adds new TDs to this landscape while permitting larger duplication populations to emerge.

These findings reveal a previously unrecognized genome-wide role for FANCM in constraining structural genome evolution and TD formation across the *BRCA1*-mutant breast cancer genome under endogenous replication stress.

### FANCM suppresses tandem duplication formation at transcriptionally active regions and limits transcription–replication conflicts

Having established that chronic FANCM depletion generates additional TDs, we next asked whether their breakpoints arise randomly or preferentially within specific chromatin environments. We examined genomic regions surrounding TD breakpoints in the baseline genome and newly emerged TDs from both FANCM-depleted clones for overlap with H3K4me3, H3K9me3 and Pol II ChIP-seq regions. Observed-to-expected ratios >1 indicate enrichment relative to random genomic expectation, whereas values <1 indicate depletion **(Fig. 4A-C)**.

**Figure 4.**
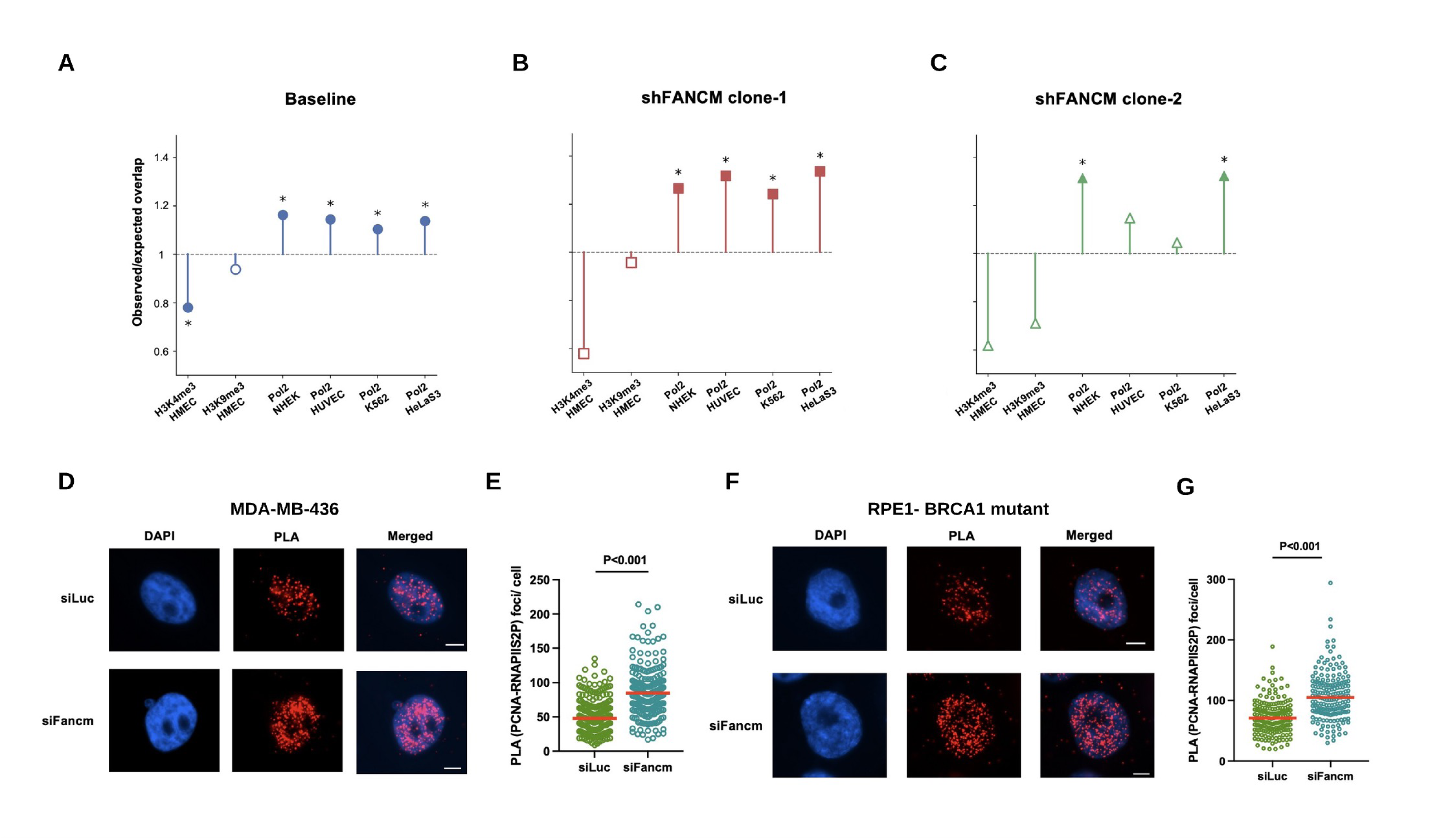
Tandem duplication breakpoint regions associate with Pol II occupancy and FANCM limits replication–transcription encounters. **a–c** Chromatin-feature enrichment analysis of genomic regions surrounding tandem duplication breakpoints in the baseline MDA-MB-436 genome (a), and newly emerged TDs in shFANCM Clone 1 **(b)**, and shFANCM Clone 2 **(c)**. Plots show observed-to-expected overlap ratios for H3K4me3 and H3K9me3 ChIP-seq regions in HMEC cells and RNA polymerase II (Pol II) ChIP-seq regions from NHEK, HUVEC, K562 and HeLa-S3 cells. The dashed horizontal line indicates an observed-to-expected ratio of 1. Filled symbols with asterisks denote statistically significant enrichment or depletion determined by permutation testing; open symbols indicate non-significant associations. **d-e** Proximity ligation assay (PLA) between PCNA and Ser2-phosphorylated RNA polymerase II (RNAPII-S2P) in *BRCA1*-mutant MDA-MB-436 cells following treatment with control siRNA (siLuc) or FANCM-targeting siRNA (siFANCM); **d**, representative images showing DAPI-stained nuclei (blue), PCNA–RNAPII-S2P PLA foci (red) and merged channels (scale bars: 5 µm); **e**, quantification of PCNA–RNAPII-S2P PLA foci per cell. **f-g,** PCNA–RNAPII-S2P PLA in *BRCA1*-mutant RPE1 cells following treatment with siLuc or siFANCM; **f**, representative DAPI, PLA and merged images and **g**, quantification of PCNA–RNAPII-S2P PLA foci per cell; for **e** and **g**, each point represents an individual cell and red horizontal lines indicate the mean. The *P* values (two-tailed Mann–Whitney test) are indicated at the top.

Baseline TD breakpoints were significantly enriched across all four independent Pol II datasets, with observed-to-expected ratios of approximately 1.11-1.16 **(Fig. 4A)**. In contrast, H3K4me3 was significantly depleted, while H3K9me3 showed a modest, non-significant depletion. Thus, Pol II occupancy, rather than generalized enrichment for individual histone marks, represented the most reproducible feature of the baseline TD breakpoint landscape.

This association was maintained and, in several comparisons, increased among TDs newly emerging after FANCM depletion. Clone 1 TD breakpoints were significantly enriched across all four Pol II datasets, with observed-to-expected ratios of approximately 1.24-1.34 **(Fig. 4B)**. Clone 2 showed a similar pattern, with significant enrichment at Pol II-bound regions in the NHEK and HeLa-S3 datasets and enrichment trends in HUVEC and K562 **(Fig. 4C)**. In contrast, H3K4me3 and H3K9me3 showed no corresponding enrichment. The reproducible association of newly emerged TDs with Pol II occupancy suggested that transcriptionally engaged genomic regions may represent preferential sites of structural instability when FANCM function is compromised.

We therefore asked whether FANCM depletion directly increases encounters between the replication and transcription machineries. Transcription–replication conflicts were assessed by proximity ligation assay (PLA) between PCNA, a component of the replication machinery, and Ser2-phosphorylated RNA polymerase II (RNAPII-S2P), which marks transcriptionally elongating RNA polymerase II. FANCM depletion in *BRCA1*-mutant MDA-MB-436 cells produced a significant increase in PCNA– RNAPII-S2P PLA foci compared with control cells **(Fig. 4D,E)**, demonstrating increased proximity between active replication and elongating transcription complexes following FANCM loss.

To determine whether this FANCM-dependent restraint of replication–transcription encounters were conserved beyond the cancer-cell context, we performed the same analysis in non-transformed RPE1 cells carrying BRCA1 deletion. FANCM depletion similarly produced a significant increase in PCNA– RNAPII-S2P PLA foci in *BRCA1*-mutant RPE1 cells compared with control cells **(Fig. 4F,G)**. Thus, increased replication–transcription encounters following FANCM loss are reproduced in genetically distinct *BRCA1*-deficient cellular backgrounds and are not restricted to the MDA-MB-436 TNBC model.

Together, the genomic and cellular analyses link FANCM-mediated TD suppression to transcriptionally active chromatin: newly emerged TDs preferentially associate with Pol II-occupied regions, while FANCM depletion increases replication–transcription encounters across cancerous and non-transformed BRCA1-deficient cells, supporting a conserved role for FANCM in limiting transcription-associated replication stress and structural genome instability.

### FANCM loss diversifies structural variant architectures in BRCA1-deficient TNBC

Having established that FANCM loss generates additional SVs and TDs, we next asked whether it alters the structural architecture of the rearrangements that emerge. FANCM depletion-induced deletions, TDs, inversions and translocations were stratified by genomic span and rearrangement configuration and compared with the pre-existing baseline landscape **(Fig. 5A-E; Supplementary Fig. 3)**.

**Figure 5.**
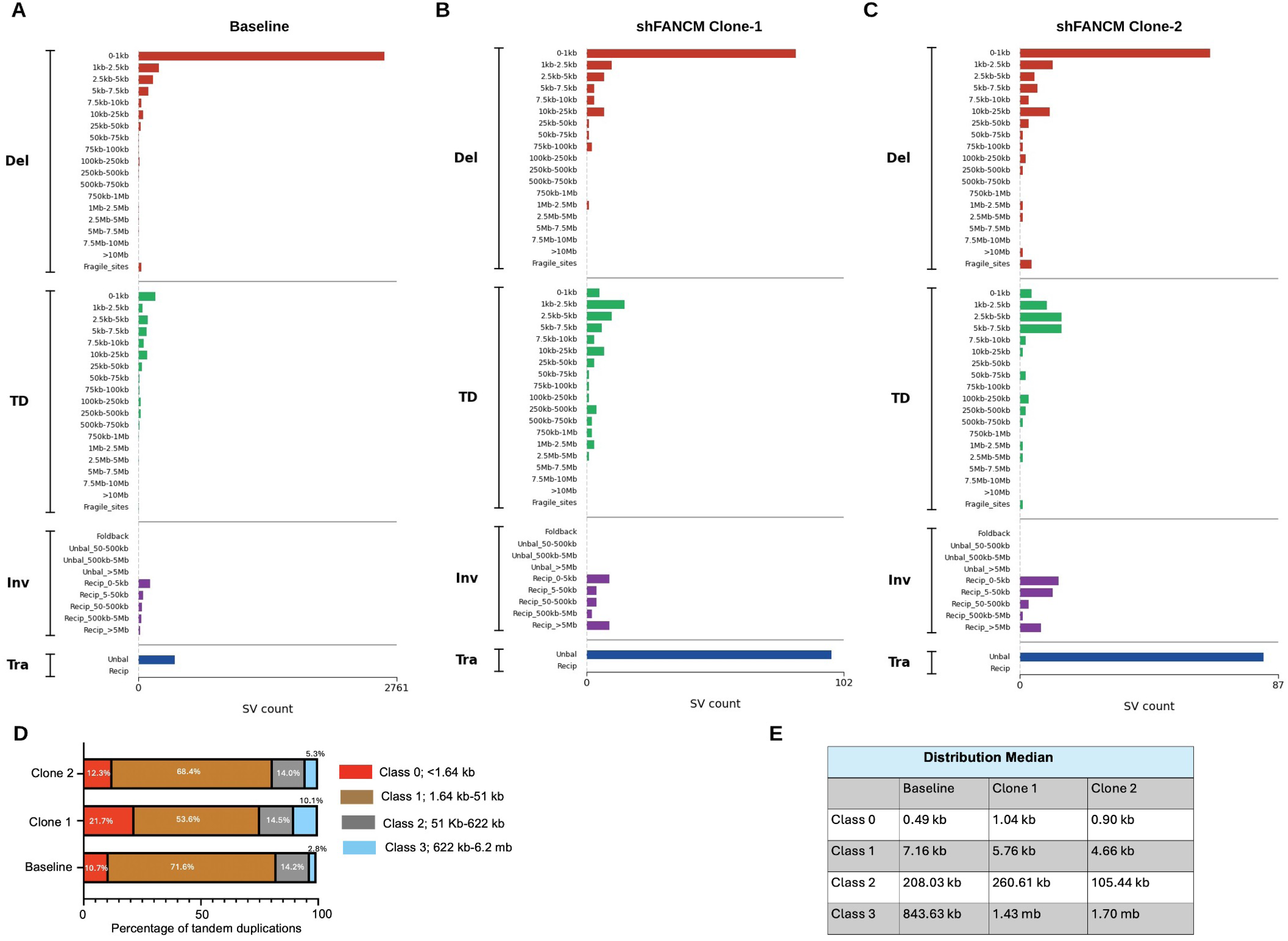
Size and structural architecture of FANCM depletion-induced genome rearrangements. **a-c**, SVs were assigned to 49 class- and size-defined categories based on the PCAWG structural variation classification framework^31^. Distribution of SVs according to class, genomic span and rearrangement configuration in the baseline *BRCA1*-mutant MDA-MB-436 genome **(a)**, and among SVs newly emerging during chronic FANCM depletion in Clone 1 **(b)** and Clone 2 **(c)**; Deletions (Del) and tandem duplications (TD) are stratified by genomic span; inversions (Inv) are classified according to configuration and span; and translocations (Tra) are classified as unbalanced or reciprocal. **d**, Percentage distribution of TDs across GMM-derived span-size classes in the baseline genome and among newly emerged TDs in Clone 1 and Clone 2. **e**, Median TD span within each GMM-derived size class for baseline, Clone 1 and Clone 2. Detailed deletion and TD size distributions are shown in Supplementary Fig. 3, and quantitative comparison of SV architectures between the independently derived FANCM-depleted clones is shown in Supplementary Fig. 4.

The baseline genome was dominated by short deletions, with 95.5% measuring <10 kb (**Supplementary Fig. 3A)**. Although <10-kb deletions remained predominant after FANCM depletion, their contribution decreased to 90.1% in Clone 1 and 76.9% in Clone 2 **(Supplementary Fig. 3B)**, accompanied by increased representation of larger events. Deletions spanning 10-100 kb accounted for 9.1% and 13.2% of newly emerged deletions in Clone 1 and Clone 2, respectively, while Clone 2 additionally accumulated events spanning 100 kb-1 Mb and >1 Mb. Thus, FANCM loss broadens deletion architecture beyond the strongly short-deletion-dominated baseline genome.

Short TDs likewise remained predominant after FANCM depletion. At baseline, 57.4% of TDs measured <10 kb, compared with 60.9% and 78.9% of newly emerged TDs in Clone 1 and Clone 2, respectively **(Supplementary Fig. 3C,D)**. GMM-derived size classes similarly showed that Class 1 TDs (1.64-51 kb) represented the largest category at baseline and in both FANCM-depleted clones **(Fig. 5D)**, indicating that combined BRCA1 and FANCM deficiency preferentially generates additional short TDs.

However, FANCM loss also diversified the TD spectrum toward substantially larger events. Clone 1 showed increased representation of Class 3 TDs spanning 622 kb-6.2 Mb (10.1% versus 2.8% at baseline), while Clone 2 likewise accumulated newly emerged Class 3 events **(Fig. 5D)**. The median span of Class 3 TDs increased from 843.63 kb at baseline to 1.43 Mb and 1.70 Mb in Clone 1 and Clone 2, respectively **(Fig. 5E)**. FANCM loss therefore produces a dual effect: amplification of the dominant short-TD phenotype together with diversification toward megabase-scale duplications.

The consequences of FANCM depletion extended beyond deletions and TDs. Reciprocal inversions emerged across multiple size ranges in both independent clones **(Fig. 5B,C)**, while newly emerged translocations were overwhelmingly represented by unbalanced rearrangements. The recurrence of these features across independent clones demonstrates that FANCM deficiency affects multiple forms of chromosome-scale structural remodeling.

To assess reproducibility across independent clones, we compared SV counts across size- and class-defined categories. Clone 1 and Clone 2 showed strong concordance in overall SV architecture (Spearman’s ρ = 0.77; Pearson’s r = 0.84; R^2^ = 0.70; **Supplementary Fig. 4A**). Class-specific analysis further revealed particularly high similarity for deletions and translocations (cosine similarity = 0.991 and 0.999, respectively), with tandem duplications and inversions also showing substantial concordance (**Supplementary Fig. 4B**). Thus, although individual rearrangements emerge independently, chronic FANCM loss generates a reproducible class- and size-specific SV architecture.

Together, these findings establish that FANCM constrains the size, diversity and structural complexity of rearrangements arising in the *BRCA1*-deficient genome.

### Chronic FANCM depletion drives genome-wide emergence of structural variants in BRCA1-deficient TNBC

We next asked whether FANCM depletion-induced SVs were restricted to particular genomic regions or emerged broadly across the *BRCA1*-deficient genome. SVs were mapped across individual chromosomes using Circos plots, with chromosome-level heatmaps providing complementary quantification of each SV class **(Fig. 6A-E)**.

**Figure 6.**
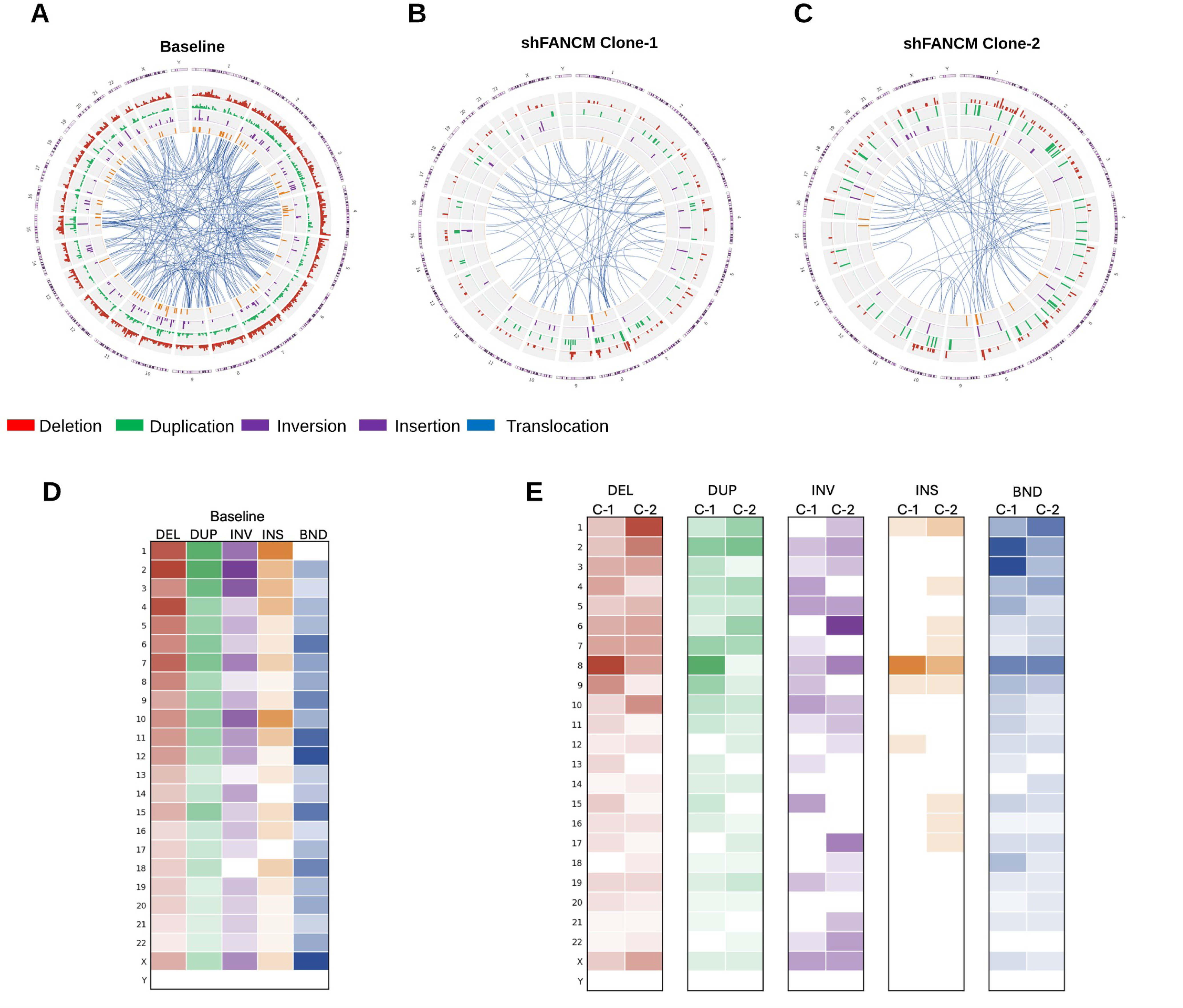
Chromosome-wide distribution of newly emerged structural variants following FANCM depletion in *BRCA1*-mutant TNBC cells. **a-c**, Circos representations of the pre-existing SV landscape of baseline *BRCA1*-mutant MDA-MB-436 cells **(a)**, and SVs newly emerging following chronic FANCM depletion in Clone 1 **(b)**, and Clone 2 **(c)**. The outer ring represents the human genome ideogram (hg38). Concentric tracks display the chromosomal density of deletions (red), duplications (green), inversions (purple), and insertions (orange), with bar height corresponding to local structural variant density. Blue links connect breakpoint pairs involved in interchromosomal translocations. **d**, Chromosome-level heatmap showing the distribution of deletion (DEL), duplication (DUP), inversion (INV), insertion (INS) and breakend/translocation (BND) events in the baseline genome. **e**, Chromosome-level heatmaps showing newly emerged SVs in Clone 1 (C-1) and Clone 2 (C-2) for each SV class. Color intensity represents relative SV abundance across chromosomes within each SV class.

The baseline MDA-MB-436 genome exhibited extensive structural variation across nearly all chromosomes **(Fig. 6A,D)**. Chronic FANCM depletion generated an additional layer of rearrangements broadly distributed across the genome in both independent clones **(Fig. 6B,C)**. Newly emerged deletions and duplications occurred across multiple chromosomes, while inversions and insertions displayed more heterogeneous chromosome-specific patterns. Both clones also contained numerous rearrangement links connecting distant genomic loci.

Chromosome-level quantification confirmed the broad distribution of newly emerged SVs **(Fig. 6E)**. Although the precise chromosomal positions and burdens differed between clones, both displayed the same overarching feature: new SVs emerged across numerous chromosomes rather than being restricted to a small number of genomic regions. Thus, FANCM deficiency produces genome-wide structural instability while permitting heterogeneity in the individual rearrangements that emerge. Together, these findings identify FANCM as a genome-wide barrier to structural rearrangement formation in BRCA1-deficient TNBC cells.

### FANCM expression stratifies structural variant architecture in *BRCA1*-altered human breast cancers

Having defined the consequences of FANCM depletion experimentally, we next asked whether FANCM expression is associated with corresponding SV architectures in human *BRCA1*-altered breast cancers. We assembled an independent cohort by integrating genomic and transcriptomic datasets from breast cancers carrying germline or somatic *BRCA1* driver alterations; tumors harboring *BRCA2* alterations were excluded. The final cohort comprised 15 *BRCA1*-altered breast cancers (13 germline and 2 somatic), which were stratified into FANCM-high and FANCM-low groups according to median FANCM expression **(Fig. 7)**.

**Figure 7.**
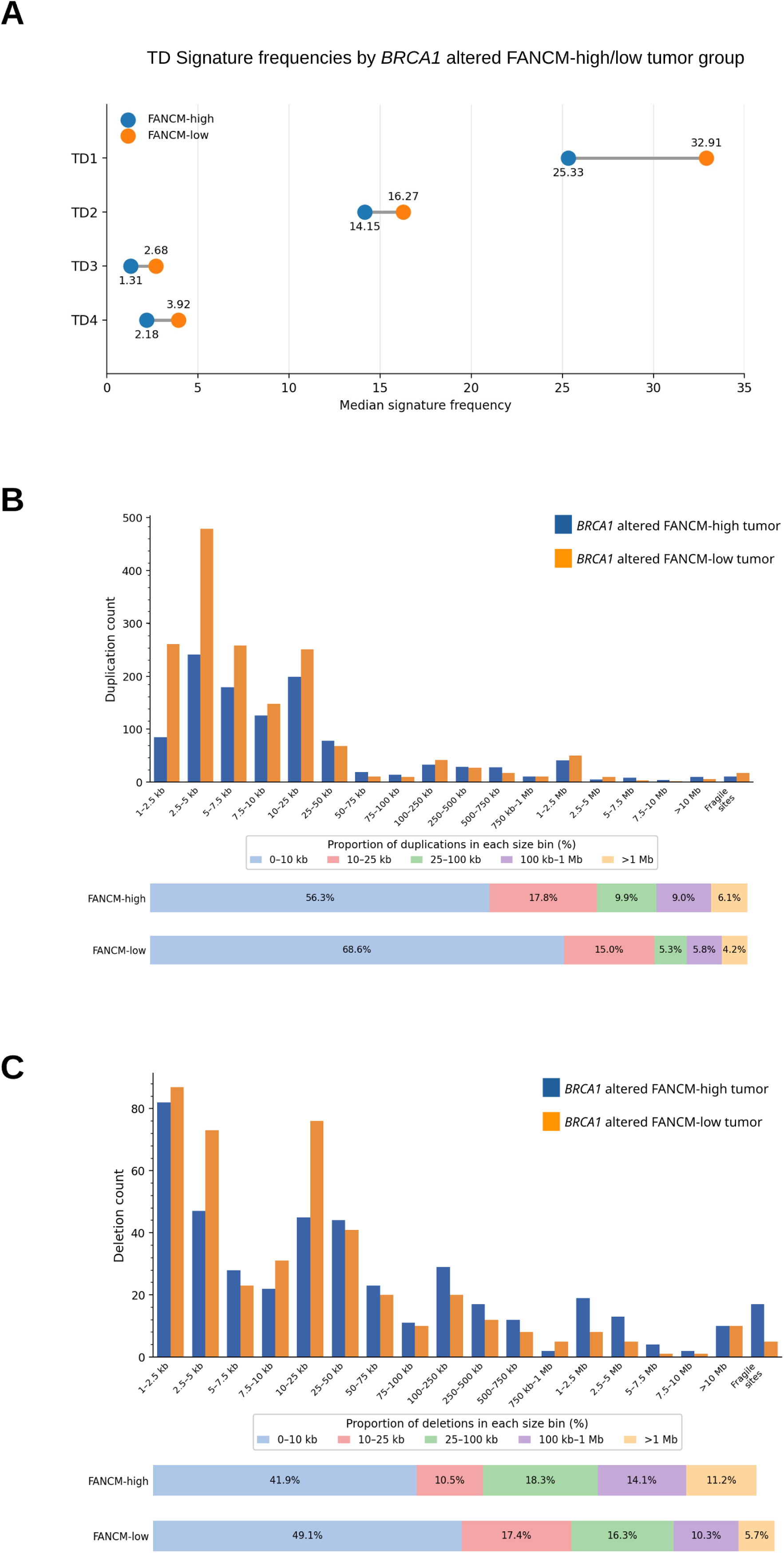
FANCM expression stratifies structural variant architecture in *BRCA1*-altered breast cancer. **a**, Median frequencies of tandem duplication signatures TD1–TD4 in FANCM-high and FANCM-low *BRCA1*-altered breast cancers. Lines connect the median signature frequencies between the two FANCM-expression groups. **b**, Size distribution of duplications in FANCM-high and FANCM-low *BRCA1*-altered tumors. Bar graph shows duplication counts across the indicated size intervals; stacked bars below show the proportion of duplications within the five aggregated size classes. **c**, Size distribution of deletions in FANCM-high and FANCM-low *BRCA1*-altered tumors. Bar graph shows deletion counts across the indicated size intervals; stacked bars below show the proportion of deletions within the five aggregated size classes.

We first examined previously defined TD signatures. FANCM-low *BRCA1*-altered tumors showed higher median frequencies across all four TD signatures than FANCM-high tumors **(Fig. 7A)**. The largest absolute difference involved the predominant TD1 signature, which increased from 25.33 in FANCM-high tumors to 32.91 in FANCM-low tumors. TD2, TD3 and TD4 were likewise higher in the FANCM-low group (14.15 to 16.27, 1.31 to 2.68 and 2.18 to 3.92, respectively). This pattern closely parallels our experimental FANCM-depletion phenotype, in which FANCM loss amplified the dominant *BRCA1*-associated short-TD phenotype while simultaneously permitting diversification toward larger, including megabase-scale, TDs **(Figs. 3 and 5)**. Thus, reduced FANCM expression in human *BRCA1*-altered tumors is associated with increased representation of multiple TD signatures, providing independent support for FANCM as a determinant of the BRCA1-associated duplication landscape.

Analysis of duplication sizes further reinforced this relationship **(Fig. 7B)**. FANCM-low tumors contained substantially greater absolute numbers of short duplications, with prominent increases across multiple kilobase-scale bins. Importantly, this increase was not restricted to events <10 kb: 10–25-kb duplications were also more numerous in FANCM-low than FANCM-high tumors, indicating expansion of the duplication burden into intermediate kilobase-scale events. Consistent with the count-based distribution, <10-kb duplications constituted 68.6% of all duplications in FANCM-low tumors compared with 56.3% in FANCM-high tumors. These human tumor data therefore recapitulate the preferential generation of short TDs observed following experimental FANCM depletion while demonstrating that the FANCM-associated duplication phenotype extends beyond the smallest size bins.

FANCM expression was also associated with deletion architecture **(Fig. 7C)**. FANCM-low tumors showed greater proportional representation of deletions measuring <10 kb (49.1% versus 41.9%) and 10–100 kb (33.7% versus 28.8%), whereas FANCM-high tumors contained proportionally more larger deletions. Thus, the relationship between FANCM expression and SV architecture in BRCA1-altered human tumors extends beyond TDs to the deletion landscape.

Together, these human tumor data independently recapitulate key features of the experimental FANCM-depletion phenotype, linking reduced FANCM expression in *BRCA1*-altered breast cancer to increased TD signatures and a pronounced burden of short-to-intermediate kilobase-scale duplications.

### Chronic FANCM depletion induces adaptive transcriptional reprogramming in BRCA1-deficient TNBC cells

Having established the structural consequences of FANCM loss and their concordance with human BRCA1-altered tumors, we next asked how cells persisting under prolonged FANCM deficiency transcriptionally adapt. RNA-seq was performed on cultures propagated in parallel with those used for WGS under the same approximately 25-population-doubling interval. Differential expression analysis identified 2,442 significantly dysregulated genes, comprising 923 upregulated and 1,519 downregulated transcripts relative to undepleted cells **(Fig. 8A)**.

**Figure 8.**
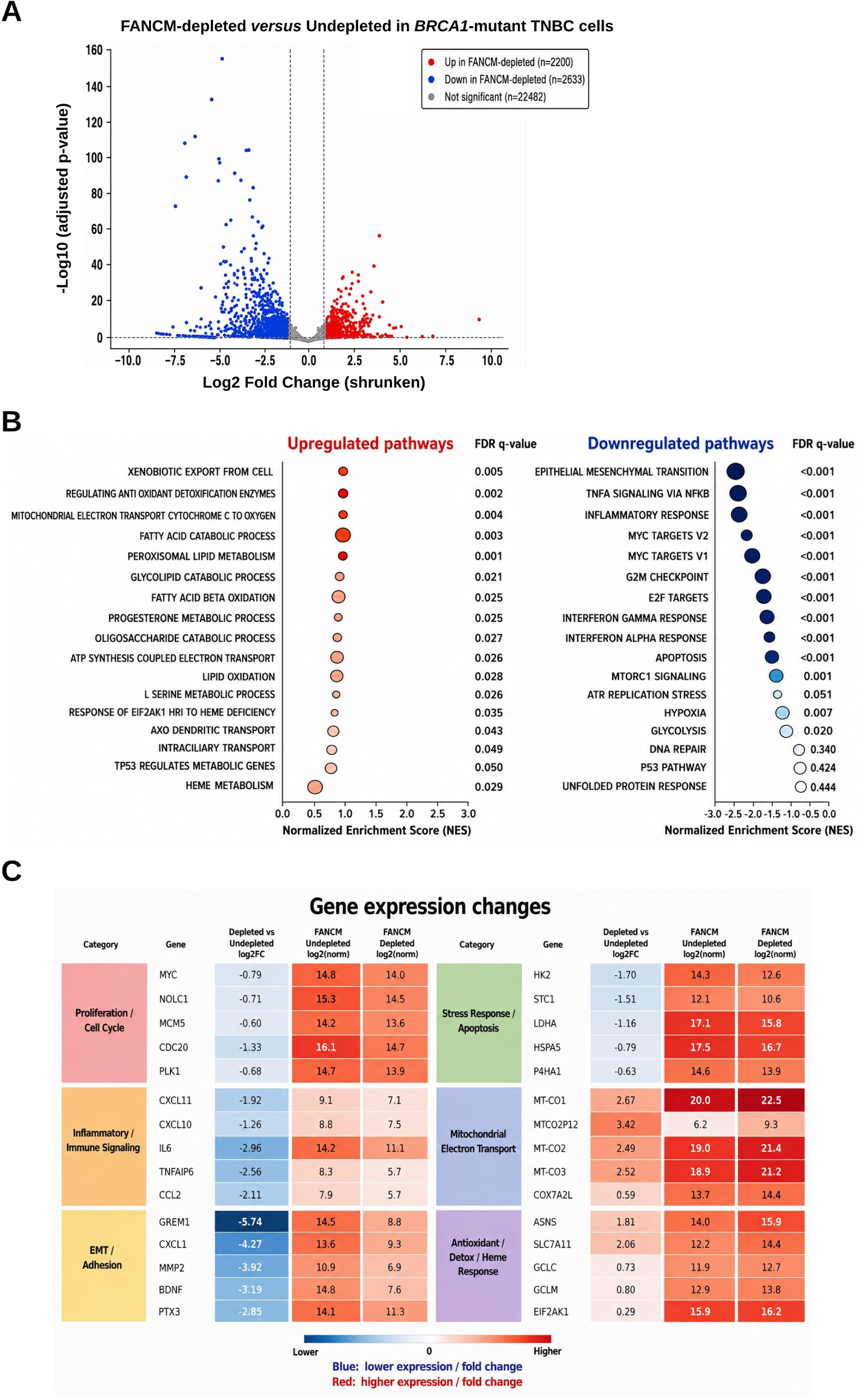
Transcriptome and pathway remodeling during chronic FANCM depletion. **a**, Volcano plot showing differential gene expression in chronically FANCM-depleted versus undepleted *BRCA1*-mutant MDA-MB-436 cells. Red and blue points denote significantly upregulated and downregulated genes, respectively; grey points denote genes not meeting the differential-expression criteria. **b**, Gene set enrichment analysis (GSEA) of pathways altered following chronic FANCM depletion. Positively enriched pathways are shown to the left and negatively enriched pathways to the right according to normalized enrichment score (NES). Symbol size represents gene-set size; FDR q values indicate enrichment significance. **c**, Heatmap of representative genes contributing to proliferation/cell-cycle, inflammatory/immune, EMT/adhesion, stress-response/apoptosis, mitochondrial electron-transport and antioxidant/detoxification programs. The left column shows FANCM depletion-associated log2 fold change, and the remaining columns show normalized expression in undepleted and FANCM-depleted cells.

Gene set enrichment analysis (GSEA) revealed coordinated suppression of proliferative programs **(Fig. 8B)**. MYC targets, E2F targets and the G2M checkpoint were strongly negatively enriched (FDR q < 0.001), while mTORC1 signaling was also significantly reduced. These changes are concordant with the progressive loss of proliferative fitness observed in the clonogenic, Incucyte and spheroid assays **(Fig. 2)**.

Inflammatory and immune-associated programs were likewise negatively enriched, including TNFα/NF-κB signaling, inflammatory response and interferon-α and interferon-γ responses **(Fig. 8B)**. Epithelial-mesenchymal transition and apoptosis were also negatively enriched, indicating broad remodeling of proliferative, inflammatory and cell-state programs. Conversely, positively enriched pathways were dominated by metabolic, mitochondrial and detoxification programs, including mitochondrial electron transport, fatty-acid catabolism and β-oxidation, peroxisomal lipid metabolism, ATP synthesis-coupled electron transport, antioxidant detoxification and heme metabolism **(Fig. 8B)**. These findings indicate a marked shift toward metabolic and stress-adaptive programs during chronic FANCM depletion.

Notably, canonical DNA damage-response transcriptional programs were not dominant at this late endpoint. ATR replication-stress signaling approached but did not meet the FDR threshold (q = 0.051), whereas Hallmark DNA repair and p53 pathways were not significantly enriched **(Fig. 8B)**. Thus, the transcriptional state of cells persisting after prolonged FANCM depletion is characterized less by a sustained canonical DNA damage response than by suppression of proliferative programs and engagement of metabolic adaptation.

Gene-level analysis supported these pathway-level changes **(Fig. 8C)**. MYC, NOLC1, MCM5, CDC20 and PLK1 were reduced, consistent with suppression of proliferative programs, while inflammatory and extracellular-remodeling genes including CXCL11, CXCL10, IL6, GREM1, CXCL1 and MMP2 were downregulated. Conversely, mitochondrial electron-transport genes and metabolic/redox regulators including ASNS, SLC7A11, GCLC and GCLM were increased.

Together, these findings reveal that chronic FANCM depletion shifts BRCA1-deficient TNBC cells from proliferative programs toward metabolic and stress adaptation, defining the transcriptional state accompanying progressive structural genome instability.

### Newly emerged structural variants are associated with local transcriptional perturbation following chronic FANCM depletion

Finally, we asked whether newly emerged SVs themselves were associated with local changes in gene expression. Integration of WGS and RNA-seq identified 503 genes intersected by or located proximal to newly emerged SVs, of which 91 were differentially expressed **(Fig. 9A)**. The remaining 412 SV-associated genes were not differentially expressed, while 4,742 differentially expressed genes lacked an identified local SV association. Thus, local SV-associated transcriptional effects represent a discrete subset of the broader transcriptional response to chronic FANCM depletion.

**Figure 9.**
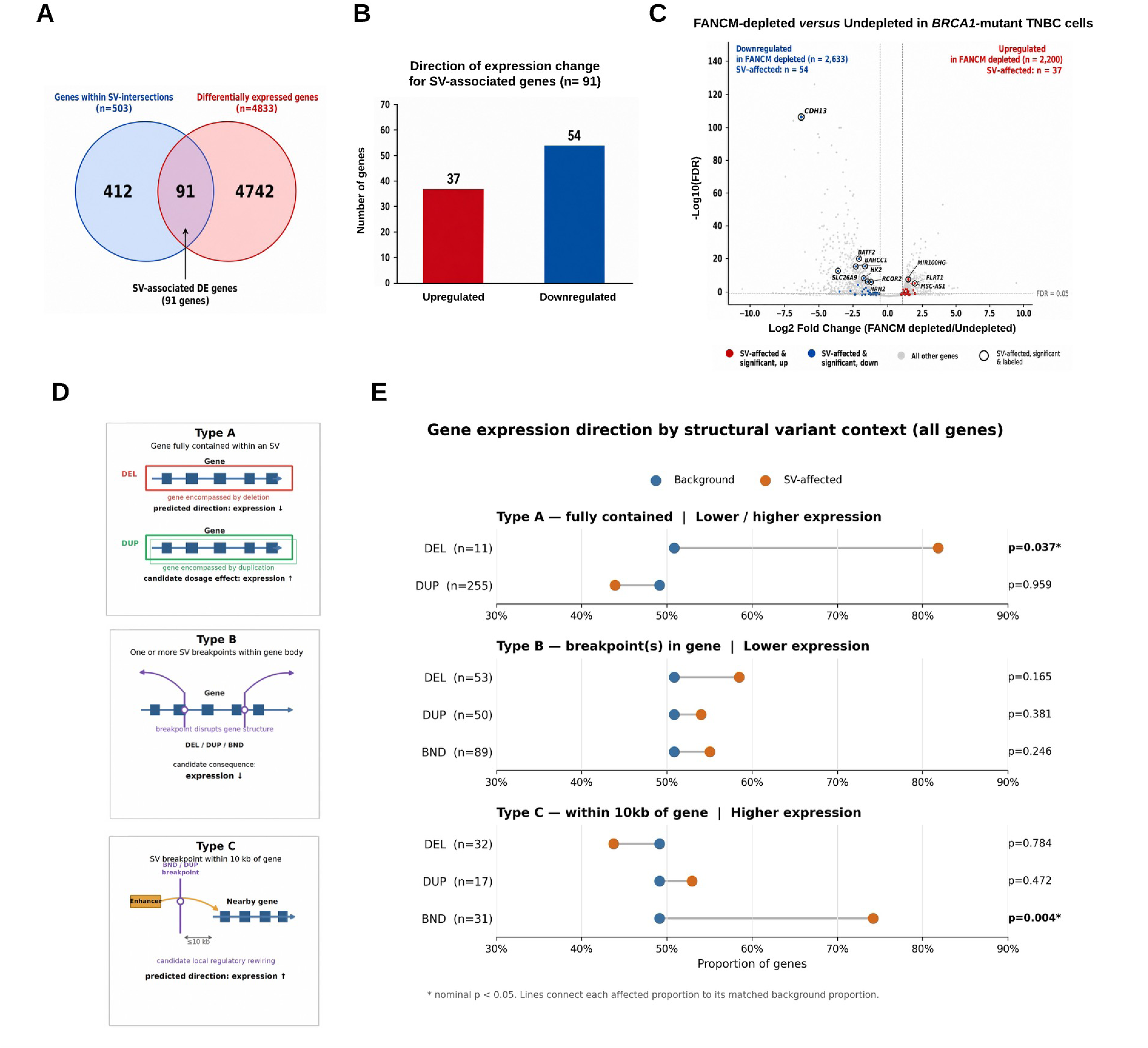
Integration of newly emerged structural variants with local gene-expression changes. **a**, Intersection of genes associated with newly emerged SVs and differentially expressed genes following chronic FANCM depletion. **b**, Direction of expression change among the 91 SV-associated differentially expressed genes. **c**, Volcano plot positioning SV-associated genes within the genome-wide transcriptional response to FANCM depletion. Significant SV-associated genes are highlighted and selected genes are labeled. **d**, Schematic defining the SV-gene relationships used for integrated WGS-RNA-seq analysis: Type A, gene fully contained within an SV; Type B, one or more SV breakpoints within the gene body; and Type C, SV breakpoint located within 10 kb of a gene. **e**, Directional expression analysis of SV-associated genes compared with SV-unaffected background genes. Type A evaluates genes fully contained within deletions (DEL) or duplications (DUP); Type B evaluates DEL, DUP or BND breakpoints within gene bodies; and Type C evaluates genes located within 10 kb of DEL, DUP or BND breakpoints. Lines connect each SV-associated proportion with its corresponding SV-unaffected background. Nominal p-values are shown; asterisks indicate significant *p-value* < 0.05.

Among the 91 SV-associated differentially expressed genes, 54 were downregulated and 37 were upregulated **(Fig. 9B)**. Mapping these genes onto the global differential-expression landscape showed that SV-associated transcripts occupied both arms of the volcano plot and spanned a broad range of effect sizes **(Fig. 9C)**.

To determine whether specific SV configurations were associated with predictable transcriptional consequences, SV-gene relationships were classified into three contexts **(Fig. 9D)**: Type A events comprised genes fully contained within an SV, Type B events contained one or more SV breakpoints within the gene body, and Type C events comprised genes located within 10 kb of an SV breakpoint. For each analysis, SV-associated genes were compared with SV-unaffected background genes, thereby distinguishing local SV-associated effects from transcriptional changes occurring independently of detectable SVs.

Among Type A events, genes fully encompassed by deletions showed the strongest association. 81.8% of 11 deletion-associated genes were downregulated compared with 50.9% of SV-unaffected genes (p = 0.037; **Fig. 9E**), representing an approximately 31-percentage-point enrichment in transcriptional loss. In contrast, genes fully contained within duplications were not significantly enriched for increased expression.

Type B events showed no significant directional enrichment. Genes intersected by deletion, duplication or BND breakpoints showed only modest increases in the frequency of downregulation relative to SV-unaffected genes, indicating that an intragenic breakpoint alone is insufficient to predict transcriptional repression.

A second significant relationship emerged among Type C events. 74.2% of genes located within 10 kb of a BND showed increased expression compared with 49.1% of SV-unaffected genes (p = 0.004; **Fig. 9E**), representing an approximately 25-percentage-point enrichment in transcriptional activation. Proximity to deletions or duplications did not produce a comparable effect. These BND-proximal events therefore identify candidate loci at which newly emerged rearrangements alter the local regulatory environment, although breakpoint proximity alone does not establish enhancer or promoter hijacking.

Together, these findings demonstrate that FANCM depletion-induced structural variants can produce configuration-dependent local transcriptional perturbations that are distinguishable from the broader SV-independent transcriptional response.

## Discussion

This study establishes FANCM as a genome-wide determinant of structural genome evolution in human BRCA1-deficient breast cancer, advancing beyond its previously defined role in repair at a single engineered replication barrier in *Brca1*-deficient mouse embryonic stem cells. By temporally exploiting the interval between FANCM depletion and synthetic lethality, we followed the consequences of FANCM loss over successive cell divisions under endogenous replication stress. Chronic FANCM depletion generated reproducible genome-wide structural variation, amplified the characteristic BRCA1-associated short-TD phenotype while permitting substantially larger TDs to emerge, and diversified additional SV classes. Newly emerged TDs preferentially associated with Pol II-occupied regions, while FANCM depletion increased PCNA–RNAPII-S2P proximity in BRCA1-mutant cancer and non-transformed cells, linking FANCM-mediated genome protection to transcription–replication encounters. Importantly, BRCA1-altered human breast cancers with low FANCM expression showed corresponding increases in TD signatures and short-to-intermediate duplications. Together, these findings establish that FANCM not only determines survival of BRCA1-deficient cells but continuously restrains structural genome evolution arising under endogenous replication stress.

The increased engagement of FANCM at nascent DNA provides a functional basis for this dependency. FANCM remodels replication intermediates, protects stalled forks and regulates repair pathway choice following replication stress^16,22^. BRCA1 itself contributes to protection of stalled replication forks, providing a mechanistic basis for increased reliance on compensatory fork-maintenance pathways when BRCA1 function is compromised^21^. BRCA1-deficient cells showed greater FANCM association with nascent DNA under basal conditions, and experimentally induced replication stress further increased FANCM fork engagement within the same *BRCA1*-mutant background. Restoration of wild-type *BRCA1* rescued sensitivity to FANCM depletion, establishing FANCM as a synthetic lethal dependency in human BRCA1-deficient TNBC. Importantly, synthetic lethality serves here not simply as an endpoint but as a temporal constraint that we exploited experimentally to interrogate what happens to the surviving cancer genome before FANCM loss becomes incompatible with continued proliferation. The requirement for FANCM across clonogenic, monolayer and three-dimensional spheroid growth further supports a model in which BRCA1 loss creates dependence on FANCM-mediated fork maintenance to sustain proliferative fitness **(Fig. 10a)**.

**Figure 10.**
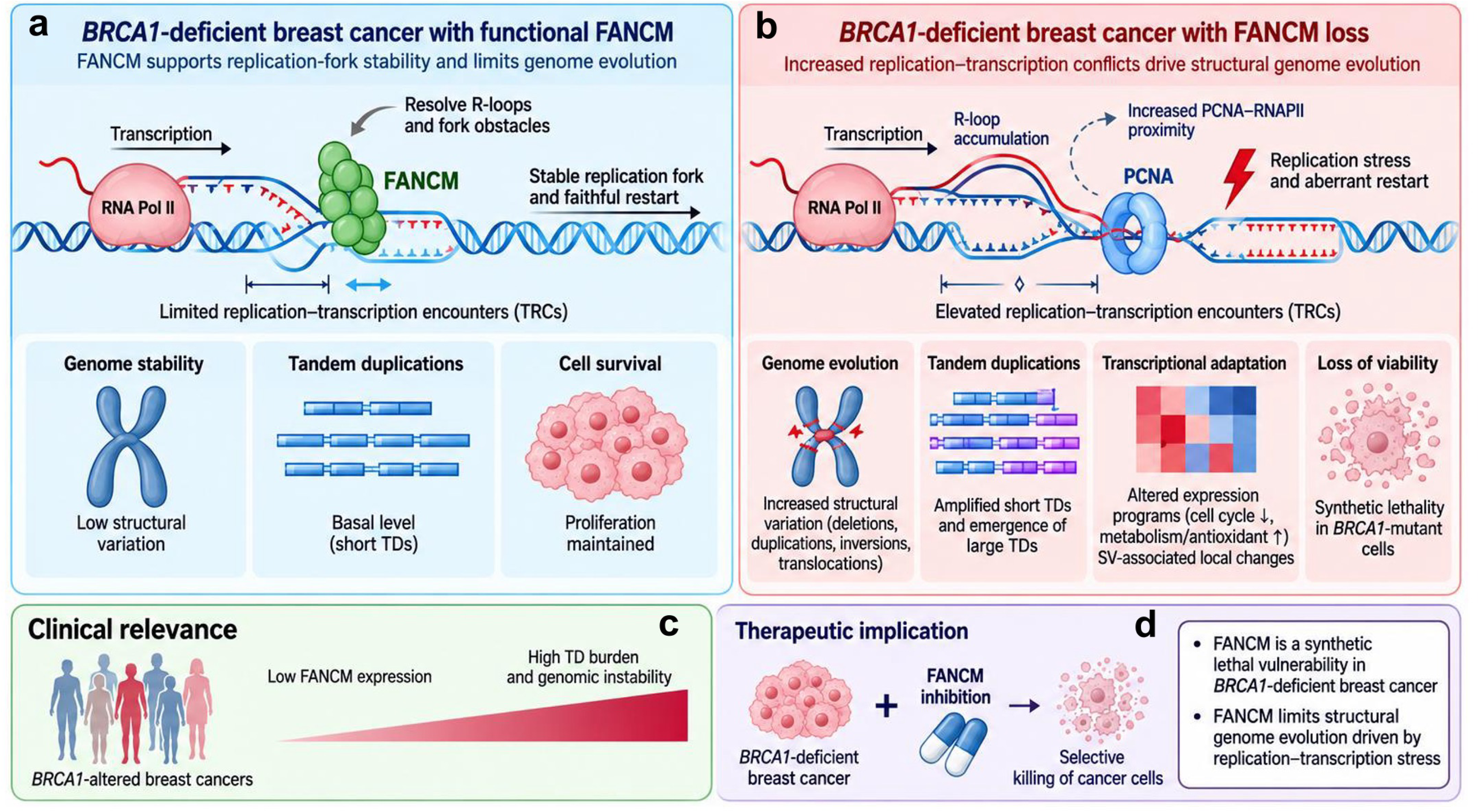
Proposed model for FANCM-dependent control of replication stress, structural genome evolution and survival in BRCA1-deficient breast cancer. **a**, functional FANCM promotes replication-fork stability, limits R-loop-associated replication–transcription encounters and restrains structural genome variation while supporting proliferation of *BRCA1*-deficient cells. **b**, FANCM loss is associated with increased replication–transcription encounters, replication stress and aberrant fork restart, accompanied by genome-wide structural variation, amplification of short TDs with emergence of larger TDs, transcriptional adaptation and progressive loss of cellular fitness. **c**, clinical association between low FANCM expression and increased TD burden and genomic instability in *BRCA1*-altered breast cancers. **d**, model depicting FANCM as a synthetic lethal vulnerability in *BRCA1*-deficient breast cancer and the potential therapeutic implication of FANCM targeting.

Our findings identify a potential endogenous context for the site-specific FANCM-dependent TD suppression previously observed at the engineered Tus/*Ter* barrier. Whereas Tus/*Ter* imposes a predefined replication block, transcriptionally active regions represent naturally occurring sources of replication stress across the cancer genome. *BRCA1*-mutant cancers are particularly susceptible to excessive TRCs, which can promote R-loop accumulation, replication stress and loss of genome integrity^28,29^. FANCM is mechanistically positioned to counter this stress: its branchpoint translocation activity can directly displace co-transcriptional R-loops *in vitro*, while FANCM deficiency increases R-loop accumulation in cells^11,17,32^. In our study, TD breakpoint-associated regions were reproducibly enriched for Pol II occupancy, including among newly emerged TDs, and FANCM depletion increased PCNA–RNAPII-S2P proximity in both cancerous and non-transformed BRCA1-mutant cells. These observations are consistent with recent genome-wide evidence linking transcription–replication collisions to TD formation in cancer^31^ and suggest that transcriptionally engaged regions represent endogenous sources of replication stress at which FANCM becomes particularly important in the BRCA1-deficient state. Together, these findings support a model in which FANCM limits transcription-associated replication stress in BRCA1-deficient cells, whereas FANCM loss increases replication– transcription encounters and the probability that perturbed forks enter rearrangement-prone restart pathways **(Fig. 10a,b)**.

Genome-wide analysis further reveals that FANCM does not simply suppress TD number; it shapes the architecture of the BRCA1-associated TD phenotype. Short TDs are a characteristic structural feature of BRCA1-deficient cancers, and previous studies at the engineered Tus/*Ter* barrier established that BRCA1 suppresses TD formation specifically at stalled replication forks^8,9^. In contrast to these site-specific observations, our longitudinal genome-wide analysis under endogenous replication stress showed that FANCM depletion produced a dual TD phenotype: preferential generation of additional short TDs, thereby amplifying the canonical BRCA1-associated architecture, together with diversification toward substantially larger, including megabase-scale, TDs. This size diversification suggests that FANCM influences not only the frequency with which TD-generating intermediates arise but potentially also the extent of the replicative process that generates the duplicated segment. BRCA1-associated TDs can arise through aberrant replication restart followed by end joining, providing a framework in which a duplicated synthesis intermediate is generated before its fixation as a chromosomal rearrangement^8,33^. FANCM is positioned upstream in this model: by remodeling perturbed forks and antagonizing mutagenic restart pathways, FANCM can restrict formation of the nascent duplication intermediate, whereas BRCA1-dependent processing provides a downstream barrier to its fixation as a TD^8,16^. Thus, the amplification and size diversification revealed by our genome-wide analysis broaden FANCM function from site-specific TD suppression to regulation of the architecture of TD formation across the BRCA1-deficient cancer genome.

The emergence of both additional short TDs and substantially larger TDs suggests a broader role for FANCM in TD-generating fork restart than previously inferred from site-specific assays. We propose that FANCM may constrain two features of this process: entry into aberrant fork restart and the extent of restart-associated DNA synthesis once initiated **(Fig. 10b)**. Increased entry of transcription-associated perturbed forks into aberrant restart following FANCM loss would increase the frequency of nascent duplication intermediates, providing a potential explanation for amplification of the short-TD phenotype. Replication-restart models of BRCA1-deficient genome rearrangement provide independent support for the idea that aberrant restarted synthesis can generate duplicated DNA segments and contribute to structurally distinct rearrangement outcomes^10,33^. FANCM-mediated remodeling or disruption of aberrant restart intermediates could therefore additionally limit their persistence and the distance over which synthesis proceeds. In the absence of FANCM, prolonged restart-associated synthesis could generate longer duplicated tracts before resolution, providing a mechanistic explanation for the emergence of larger, including megabase-scale, TDs. FANCM loss could therefore exert two separable effects on the BRCA1-associated TD phenotype—increasing the frequency of TD-generating intermediates while broadening their potential span. Because restart-associated synthesis was not directly measured here, regulation of synthesis-tract length by FANCM remains a mechanistic hypothesis; however, the reproducibility of the size-diversified TD phenotype across independently evolving clones, together with its concordance with FANCM-low BRCA1-altered human tumors, provides a strong rationale for testing whether FANCM controls both initiation and progression of TD-generating fork restart.

This model provides a mechanistic framework for understanding why combined BRCA1 and FANCM deficiency is particularly destabilizing **(Fig. 10b,c)**. Rather than acting redundantly, FANCM and BRCA1 appear to provide complementary barriers at different stages of TD formation. FANCM acts upstream by limiting persistent fork perturbation and entry into aberrant restart pathways, thereby restricting the generation and potentially the span of nascent duplication intermediates, whereas BRCA1-dependent DNA-end processing acts downstream to prevent their fixation as stable TDs^8,16,33^. Consequently, FANCM loss in a BRCA1-deficient background couples increased generation of TD-forming intermediates with impaired capacity for their conservative resolution, providing a mechanistic basis for both the synthetic lethal interaction and the pronounced structural genome instability observed when both protective pathways are compromised. This sequential loss of replication-fork safeguards therefore links BRCA1–FANCM synthetic lethality to continued structural genome evolution rather than treating these phenotypes as independent consequences of combined deficiency.

The relevance of this genome-wide FANCM function is strengthened by its convergence with human breast cancer genomes. BRCA1 deficiency is strongly associated with the short-span TD phenotype^9,9^ and within our independent cohort of *BRCA1*-altered breast cancers, lower FANCM expression was associated with increased representation of multiple TD signatures, with the strongest absolute difference involving the predominant TD1 signature, together with a greater burden of short-to-intermediate duplications. This pattern parallels the experimentally induced FANCM-loss phenotype, in which the dominant BRCA1-associated short-TD architecture was amplified while additional TD size classes emerged. Although FANCM expression is an associative rather than functional measure of FANCM activity in these tumors, the concordance between temporally controlled FANCM depletion and independently derived human tumor genomes suggests that variation in FANCM activity may contribute to heterogeneity of the BRCA1-associated duplication landscape in human disease. Thus, the genome-wide structural phenotype uncovered experimentally is not restricted to a cell-culture model but is reflected in the genomic architecture of BRCA1-altered human breast cancers **(Fig. 10c)**.

Importantly, FANCM-mediated genome protection extends beyond TD suppression to multiple classes of structural variation across the BRCA1-deficient genome. Chronic FANCM depletion generated deletions spanning broader size ranges, reciprocal inversions and predominantly unbalanced translocations, with newly emerged rearrangements distributed across numerous chromosomes. Despite differences in individual breakpoint positions, independently evolving FANCM-depleted clones showed strong concordance in their class- and size-specific SV architectures. This reproducibility suggests that FANCM loss does not simply increase stochastic genome damage or specify particular rearrangements but instead biases the replication-stressed BRCA1-deficient genome toward a characteristic spectrum of structural outcomes. Thus, FANCM functions as a genome-wide barrier to structural diversification under endogenous replication stress, substantially broadening its previously defined role in suppressing TD formation at an individual engineered stalled fork.

Structural genome evolution following chronic FANCM loss was accompanied by substantial remodeling of the transcriptional state of surviving BRCA1-deficient cells. Cells persisting through prolonged FANCM depletion suppressed proliferative, cell-cycle and inflammatory programs while enriching mitochondrial, metabolic and antioxidant pathways, consistent with a transition toward a stress-adapted state. Notably, canonical DNA-repair and p53 transcriptional programs were not dominant at this late endpoint, suggesting that cells surviving chronic FANCM loss had progressed beyond an acute damage response toward a distinct adaptive state. These findings indicate that prolonged FANCM deficiency affects not only genome architecture but also the cellular programs that accompany survival under sustained replication stress.

Integration of WGS and RNA-seq further revealed that newly emerged SVs are associated with configuration-dependent local transcriptional consequences. Genes encompassed by deletions were preferentially downregulated, whereas genes proximal to BNDs were preferentially activated relative to SV-unaffected genes; other SV configurations showed no comparable directional association. These findings distinguish local SV-associated effects from the broader transcriptional adaptation to chronic FANCM loss and suggest that structural genome evolution can reshape the transcriptional environment encountered during subsequent rounds of DNA replication. Given the increased replication–transcription encounters following FANCM depletion, it will be important to determine whether SV-associated transcriptional activation creates or reinforces genomic regions susceptible to transcription-associated replication stress, potentially linking newly acquired rearrangements to further genome instability.

Several limitations define important directions for testing this model. Although BRCA1–FANCM synthetic lethality was observed across multiple TNBC models and FANCM loss increased replication–transcription encounters in both cancerous and non-transformed BRCA1-mutant cells, longitudinal genome evolution was examined in a single BRCA1-mutant TNBC background. Establishing causality will require directly linking R-loop accumulation, replication–transcription encounters, fork restart and subsequent TD formation, and determining whether FANCM controls the persistence or synthesis-tract length of restarted intermediates. Extending these analyses to additional BRCA1-mutant models, patient-derived organoids and in vivo systems will further establish the generality of FANCM-dependent genome protection and the therapeutic potential of targeting this synthetic lethal dependency.

In summary, our findings establish FANCM as an adaptive genome-maintenance factor that enables BRCA1-deficient cancer cells to tolerate endogenous replication stress while continuously restraining structural genome evolution. By moving beyond site-specific repair at an engineered replication barrier and temporally interrogating FANCM loss over successive cell divisions, this study reveals a genome-wide role for FANCM in limiting transcription-associated fork perturbation, TD amplification and size diversification, and broader structural rearrangement in human BRCA1-deficient breast cancer. The convergence of these experimental phenotypes with BRCA1-altered human tumor genomes further supports the disease relevance of this function. Together, these findings position FANCM at the intersection of replication-stress tolerance, structural genome evolution and synthetic lethal vulnerability in BRCA1-deficient breast cancer **(Fig. 10)**.

## Methods

### Fraction Genome Altered Relative to *BRCA* Gene Status

Publicly available datasets from three independent breast cancer cohorts were obtained via the cBioPortal platform (https://www.cbioportal.org): *TCGA PanCancer Atlas*, *MSK 2025*, and *TCGA Nature 2012*. All datasets were accessed under the “Breast” category, specifically within “*Invasive Breast Carcinoma”* studies. For each dataset, somatic mutations and putative copy-number alterations derived from GISTIC were selected for all subsequent analysis. Queries were performed using either *BRCA1* as the gene of interest, and all available samples were included. Samples were stratified into *mutant* and *wild-type* groups based on gene status. Cases lacking mutation profiling data for *BRCA1* were excluded. Fraction of Genome Altered (FGA) was extracted as a clinical attribute for each sample. Corresponding patient-level data, including patient identifiers, *BRCA1* mutation status, and FGA values, were downloaded for downstream analysis. All statistical analyses and data visualization were performed using GraphPad Prism (version 10.6.1).

### Cell Culture

MDA-MB-436, HCC-1395, MDA-MB-231 and MDA-MB-468 cell lines were purchased from ATCC. MDA-MB-436+BRCA1 (add-back) cell line was generously gifted by Neil Johnson (Fox Chase Cancer Center, Philadelphia, PA), HCC-1937 and H578T cell lines were generously gifted by John Hawse (Mayo Clinic, Rochester, MN). RPE1-hTERT TP53^−/−^ (referred to as RPE1-Wild type in this paper) and RPE1-hTERT TP53^−/−^ BRCA1^−/−^ (referred to RPE1-BRCA1 mutant in this paper) were generously gifted by Daniel Durocher (University of Toronto, Toronto, Canada). MDA-MB-436 (and its BRCA1 add-back cell line), MDA-MB-231 and MDA-MB-468 cell lines were cultured in RPMI-1640 medium supplemented with 10% fetal bovine serum and 1% Pen/Strep. All cell lines were regularly tested for mycoplasma infection by Myco-Alert assay (Lonza, LT07-318). Inducible shFANCM (HV-TRE)/shSracmbled (HV-scrambled) transduced MDA-MB-436 cells (described below) were cultured in RPMI-1640 medium supplemented with 10% fetal bovine serum (Tet system approved, Thermo Fisher Scientific, A473640) and 1% Pen/Strep. HCC-1937, HCC-1395 and H578T cell lines were cultured in 1:1 Dulbecco’s Modified Eagle Medium/Nutrient Mixture F-12 (DMEM/F-12) supplemented with 10% fetal bovine serum and 1% Pen/Strep. *RPE1*-Wild Type and RPE1-*BRCA1* mutant were cultured in DMEM/F-12 + 1% Pen/Strep + 10% fetal bovine serum + 1% L-glutamine. All cells were maintained under standard conditions (humidified atmosphere, 95% air, 5% CO2, 37 °C).

### Reverse-transcription polymerase chain reaction (RT-qPCR) analysis

RNA was extracted from cell lines and using RNeasy Mini Kit (Qiagen 74104),and from mouse TNBC PDX tumor models using Qiagen’s RNA Plus easy mini kit (Cat. #74136). FANCM, BRCA1 and reference gene expression levels were analyzed using an Applied Biosystems 7300 Real time PCR System or QuantStudio 3 using Power SYBR Green RNA-to CT ™ 1-Step Kit (Applied Biosystems, 4368702). Experiments were performed and analyzed in at least 3 independent biological triplicates. For tumor cell lines, FANCM gene expression level was normalized to GAPDH, and for TNBC PDX tumor models, FANCM and BRCA1 gene expression levels were normalized to β-actin using the 2−ΔCT method. For RT-qPCR analysis of human genes, the following primer pairs were used: FANCM, forward 5′-AAATGACAGGGTCTACACAA-3′ and reverse 5′-CTTTATTTCAGCAGCGGGA-3′; ACTB (β-actin), forward 5′-ACAGAGCCTCGCCTTTG-3′ and reverse 5′-CCTTGCACATGCCGGAG-3′; BRCA1, forward 5′-ATTCAAACTTAGGTGAAGCAGCATCT-3′ and reverse 5′-GTATCCCTCTGCTGAGTGGT-3′; and GAPDH, forward 5′-TACTAGCGGTTTTACGGGCG-3′ and reverse 5′-TCGAACAGGAGGAGCAGAGAGCGA-3′.

### Stratification by Tumor Stage and Molecular Subtype Relative to FGA

To assess the relationship between tumor stage, molecular subtype, and FGA, data from the *TCGA PanCancer Atlas* breast invasive carcinoma cohort were obtained via cBioPortal. FGA values were extracted for all available samples irrespective of *BRCA1* mutation status. Gene expression data were retrieved by querying for FANCM as gene of interest across all samples. The mRNA expression z-scores relative to diploid samples (RNA-Seq V2 RSEM) were visualized using the heatmap function within the “Tracks” module. Clinical annotations, including tumor stage and molecular subtype, were simultaneously obtained by selecting “American Joint Committee on Cancer Tumor Stage Code” and “Subtype” from the “Clinical” data tracks. Samples with low FANCM expression (z-score < 0) were excluded, and subsequent analyses were restricted to cases with high FANCM expression (z-score > 0). Corresponding FGA values for these samples were then analyzed. For downstream stratification, samples were grouped according to tumor stage (T classification) and molecular subtype. Tumor stage classification followed the TNM system based on primary tumor size: T1 (≤ 2 cm; including T1, T1a, T1b, T1c), T2 (> 2 cm to ≤ 5 cm), T3 (> 5 cm), and T4 (tumors with direct extension to the chest wall and/or skin, including inflammatory breast cancer subtypes such as T4d). Molecular subtype analyses excluded samples classified as “Normal-like.” FGA distributions were subsequently compared across tumor stage and subtype groups. Data visualization and statistical analyses were performed using GraphPad Prism (version 10.6.1), with specific tests and sample sizes detailed in the corresponding figure legends.

### HRD Score Analysis

HRD scores as a clinical attribute for 687 samples from the TCGA PanCancer (PANCAN) dataset, were extracted from UCSC Xena interface^34^ (https://xena.ucsc.edu/). Corresponding patient-level data, including patient identifiers, *BRCA1* mutation status and histological staining results were also all downloaded from Xenabrowser datahubs for interpretation and downstream analysis. Patients were then stratified based on breast cancer subtype into Basal-like, Luminal A/B or HER2 and then further stratified based on BRCA1 status for further analysis. After excluding silent mutations, we retained 11 patients with BRCA1 mutation mutations. Associations between HRD scores, cancer subtype and BRCA1 status were plotted and analyzed using GraphPad Prism (version 10.6.1).

### SiRNA/shRNA-mediated FANCM depletion

For siRNA-mediated FANCM depletion, cells were seeded at 3 × 10⁵ cells per well in 6-well plates 24 h before transfection. siRNA SMARTpools (Horizon Discovery/Dharmacon) were transfected using Lipofectamine RNAiMAX (Invitrogen, 13778150) in Opti-MEM according to the manufacturer’s instructions. siRNA–lipid complexes were incubated for 15 min at room temperature before addition to the cells. For short hairpin RNA (shRNA)-mediated targeted depletion of FANCM, a shRNA sequence (5′-GAACAGAAGTAGAAAGAAAAG-3′) specific to the FANCM transcript was cloned into the third-generation lentiviral pLKO.1-Puro expression vector (Addgene plasmid #10878). Lentiviral particles carrying either the FANCM-targeting shRNA or empty vector were generated in HEK293T packaging cells by co-transfection of the shRNA expression plasmid with the helper plasmids pX-PAX2 and pMD2 using a CaCl₂ based transfection method. Following transfection, the culture medium was replaced with fresh medium the next day. Viral containing supernatants were harvested 48 h later and concentrated 10-fold using 40% PEG-800 (w/v). HCC-1937 cells were subsequently infected with the concentrated lentiviruses containing either the empty vector or FANCM-shRNA. The culture medium was replaced with fresh medium the following day, and infected cells were subjected to puromycin selection (4µg/mL, 2 days). The efficiency of FANCM silencing was subsequently assessed by RT-qPCR. Following antibiotic selection, viable cells were collected, counted, and subsequently seeded in 6-well plates for clonogenic survival analysis.

### Generation of doxycycline inducible shRNA targeting FANCM in *BRCA1-*mutant MDA-MB-436 cell lines

A dual-vector tTS/rtTA Tet-On lentiviral system (VectorBuilder) was used to achieve inducible FANCM knockdown with minimal leaky shRNA expression. The Tet regulatory vector expresses both tTS and rtTA and carries an mCherry reporter, whereas the second vector contains the three FANCM-targeting shRNA (sequences reported below) under the control of a tetracycline response element (TRE) and carries a GFP reporter. In the absence of doxycycline, tTS suppresses TRE-driven transcription, while doxycycline enables rtTA-mediated activation. FANCM-targeting shRNA sequences were designed against the FANCM open reading frame, assessed for target specificity, and cloned into the TRE-regulated lentiviral vector. Lentiviral particles carrying either the tTS/rtTA regulatory construct or the TRE-shFANCM construct were produced in HEK293T cells. MDA-MB-436 cells were sequentially transduced with the two lentiviral vectors. Cells were first transduced with the tTS/rtTA regulatory lentivirus in the presence of polybrene and selected with hygromycin to establish a stable Tet regulatory cell population. These cells were subsequently transduced with the TRE-shFANCM lentivirus and selected with blasticidin to generate the final inducible FANCM-knockdown cell line, termed MDA-MB-436 HV-TRE (or HV-Scrambled control). In the Pool population, FANCM knockdown was induced by doxycycline (Sigma-Aldrich, Cat. # D5207) treatment and validated by RT-qPCR.

Single-cell-derived clones were generated by low-density plating followed by cloning-ring isolation. Cells were seeded sparsely to allow individual colonies to arise from single cells; well-isolated colonies were identified microscopically, enclosed using sterile cloning rings, detached with trypsin-EDTA, and transferred to individual wells for clonal expansion. In the resulting single-cell-derived clones, FANCM knockdown was induced by doxycycline (Sigma-Aldrich, Cat. # D5207) treatment and validated by RT-Qpcr

ShFANCM sequences:

shFANCM [miR30-shRNA #1]: 5′-GACTTCATGAAACTCTATAAT-3′
shFANCM [miR30-shRNA #2]: 5′-AGGACGAGAGGAACGTATTTA-3′
shFANCM [miR30-shRNA #3]: 5′-CACACCAGGTAGTGATATAAA-3′

### Clonogenic survival

Cells were seeded at low density in 6-well plates and allowed to adhere overnight. Cells were subsequently cultured for approximately 10–14 days, or until visible colonies had formed. Culture medium for DOX-treatment conditions was replaced every 3–4 days. At the experimental endpoint, colonies were washed gently with PBS, fixed with 4% paraformaldehyde, and stained with crystal violet. Excess stain was removed by washing with water, and plates were allowed to air-dry. Colonies were imaged and quantified using ImageJ (version 1.54p).

### Live Cell Imaging

MDA-MB-436 (HV-TRE or HV-Scrambled) cells were seeded into doxycycline-containing medium in 96-well tissue culture plates at a density optimized to maintain logarithmic growth throughout the duration of the experiment. Phase-contrast images were acquired at regular intervals using an IncuCyte SX5 Live-Cell Analysis System (Sartorius), and percent confluence was quantified using IncuCyte integrated analysis software. An AI-based cell recognition algorithm was used to identify cells across all time points with analyzer parameters optimized as needed to ensure accurate segmentation^35^. Cell proliferation was expressed as percent confluence over time. Data are presented as the mean ± SD of four independent wells per condition. Growth curves were analyzed by Repeated Measures two-way ANOVA using a full factorial model with treatment and time as factors. Multiple comparisons between treatment groups at individual time points were performed using the Holm–Šidák correction. A two-sided P value <0.05 was considered statistically significant.

### Spheroid Formation Assay

MDA-MB-436 HV-TRE and their scrambled control cells were grown to confluence prior to initiation of the spheroid assay. Matrigel was thawed on ice before use and maintained on ice throughout the procedure to prevent premature polymerization. Pipette tips used for handling Matrigel were pre-cooled at −80°C. Cells were trypsinized, resuspended in 1× PBS, and counted. For each condition, 4,000 cells were transferred to individual microcentrifuge tubes and centrifuged at 2,500 rpm for 5 min. The supernatant was carefully removed, and the cell pellet was resuspended in 30 µL Matrigel using pre-cooled pipette tips. The cell–Matrigel suspension was immediately transferred into wells of a 96-well plate. Plates were incubated at 37°C for 30 min to allow Matrigel polymerization, after which 100 µL of complete culture medium was added to each well. Doxycycline treatment was initiated 24 h after embedding the cells in Matrigel. Doxycycline-containing medium was prepared and added at a final concentration of 2 µg/mL. Control wells (HV-TRE and HV-Scrambled) received an equivalent volume of doxycycline-free medium. Every 3–4 days, 100 µL of medium was carefully removed from each well and replaced with 100 µL of fresh medium containing the appropriate treatment. Spheroid formation and growth were monitored until the experimental endpoint. Spheroids were imaged using a Cytation 5 Cell Imaging Multi-Mode Reader under bright-field illumination. Images were acquired using 4× and 10× objectives (for representation) at approximately days 3-12 and plates were maintained in culture for approximately 18–20 days in total.

### In situ analysis of protein interactions at DNA replication forks (SIRF) and proximity ligation assay (PLA)

SIRF and PLA were performed using the Duolink In Situ PLA system (Millipore Sigma, DU092008) according to the manufacturer’s instructions. SIRF was performed essentially as previously described^36^.For SIRF, cells were pulse-labeled with 125 µM 5-ethynyl-2′-deoxyuridine (EdU) for 2 h to label nascent DNA, followed by incubation for an additional 2 h in fresh medium or medium containing 0.5 mM hydroxyurea (HU) or 10 µM camptothecin (CPT) to induce replication stress. Click-iT chemistry was subsequently performed to conjugate biotin to EdU-labeled nascent DNA, enabling detection of proteins in proximity to newly synthesized DNA.

For both SIRF and conventional PLA, cells were fixed, permeabilized and incubated with the indicated primary antibodies overnight. For SIRF, proximity between biotin-labeled nascent DNA and FANCM was detected using antibodies against biotin and FANCM. For PLA, proximity between the replication and transcription machineries was assessed using antibodies against proliferating cell nuclear antigen (PCNA) and Ser2-phosphorylated RNA polymerase II (RNAPII-S2P). The following day, species-specific Duolink PLA probes were applied, followed by ligation and rolling-circle amplification according to the manufacturer’s protocol. Slides were mounted using DAPI-containing mounting medium. Slides were imaged using a Nikon Eclipse Ti2 microscope. SIRF and PLA foci were quantified as the number of fluorescent puncta per nucleus/cell using image J with ∼150 cells analyzed per condition. *p*-values are derived using the two-tailed Mann–Whitney test.

Primary antibodies were rabbit anti-biotin (D5A7, Cell Signaling Technology), anti-FANCM (sc-101389, Santa Cruz Biotechnology), anti-PCNA (sc-56, Santa Cruz Biotechnology) and anti-RNAPII-S2P (NB100-1805, Novus Biologicals).

### Whole-genome sequencing

Genomic DNA from the baseline MDA-MB-436 population and the two independently derived chronically FANCM-depleted clones was isolated using QIAamp DNA Micro Kit (Qiagen,56304) and subjected to whole-genome sequencing by Novogene. Whole-genome libraries were sequenced on an Illumina NovaSeq X Plus platform using 150-bp paired-end (PE150) sequencing, with a target output of 90 Gb raw sequence data per sample (∼30× nominal genome coverage) and a sequencing quality specification of Q30 ≥85%. Raw sequencing reads were processed using the open-source Sarek pipeline in tumor–normal mode^37^. The baseline MDA-MB-436 genome was used as the matched reference for identification of structural variants that emerged during chronic FANCM depletion.

### Structural variant calling, filtering and classification

Somatic structural variants were called using Manta v1.6.0. Each chronically FANCM-depleted clone was compared with the baseline reference to identify newly emerged SVs during the FANCM-depletion interval. SV calls were filtered using a minimum Manta SOMATICSCORE of 10. Variants overlapping ENCODE blacklist regions were excluded to minimize calls arising from repetitive, poorly mappable or otherwise problematic genomic regions. Following filtering, SVs were classified as: deletions (DEL), duplications (DUP), inversions (INV), insertions (INS), or breakend/translocation events (BND). SVs were assigned to 49 class- and size-defined categories based on the PCAWG structural variation classification framework^31^. For each sample or clone, the proportion represented by each aggregate size category was calculated as the number of events within that category divided by the total number of events of the corresponding SV class.

### Tandem duplication analysis

The tandem duplicator phenotype (TDP) score was calculated using the approach described by Menghi *et al*. to quantify the genome-wide distribution of tandem duplications across autosomes^5^. Samples with a TDP score >0 were considered to exhibit evidence of a tandem duplicator phenotype and were subjected to further TD size-distribution analysis. Gaussian mixture models (GMMs) were fitted to the log10-transformed TD span distributions to identify underlying TD size populations. TD size classes were interpreted with reference to the characteristic TD distributions described by Menghi *et al.*, including approximately: Class 1: short TDs centered near ∼11 kb, Class 2: intermediate TDs centered near ∼231 kb, Class 3: large TDs centered near ∼1.7 Mb^9^. The experimentally observed distributions were not forced to these modal values; rather, GMM-derived modes were estimated from each dataset and then interpreted in relation to established TD classes.

### Reproducibility of structural variant architecture between independent clones

To determine whether chronic FANCM depletion generated reproducible SV architectures across independently evolving clones, SV counts from Clone 1 and Clone 2 were compared across the 49 class- and size-defined SV categories.

Categories containing events were compared using both rank- and linear-correlation metrics. Spearman’s rank correlation coefficient (ρ) was used to assess monotonic concordance between clones, while Pearson’s correlation coefficient (r) quantified linear concordance. The coefficient of determination (R^2^) was additionally calculated from the clone-to-clone comparison.

Class-specific similarity was independently assessed using cosine similarity and Jensen–Shannon divergence (JSD). Cosine similarity measures similarity in the direction of the multidimensional SV-category vectors, with values approaching 1 indicating highly similar distributions. JSD measures divergence between normalized distributions, with values approaching 0 indicating increasing similarity. These metrics were calculated separately for deletions, tandem duplications, inversions and translocations. The resulting values were used to quantify reproducibility of class-specific SV architecture between the two independently derived FANCM-depleted clones

### Chromatin enrichment analysis at tandem duplication breakpoints

To determine whether TD breakpoints preferentially localized to specific chromatin environments, genomic regions surrounding TD breakpoints were tested for overlap with publicly available ENCODE ChIP-seq peak datasets. Analyses were performed independently for pre-existing TDs in the baseline sample and TDs newly emerged following chronic FANCM depletion in Clone 1 and Clone 2.

ChIP-seq peak calls were obtained as BED files from the UCSC Genome Browser ENCODE Broad Histone repository (hg19; wgEncodeBroadHistone). Six datasets representing transcriptionally active and repressive chromatin environments were analyzed, comprising H3K4me3 and H3K9me3 in HMEC, together with RNA polymerase II (Pol II) occupancy in NHEK, HUVEC, K562 and HeLa-S3 cells. Because the downloaded ENCODE peak files were provided in hg19 coordinates, genomic coordinates were converted to GRCh38/hg38 using UCSC LiftOver before overlap analysis.

Following the breakpoint-centered approach used by Menghi *et al.*, each TD breakpoint was extended by 200 kb upstream and 200 kb downstream, generating a 400-kb breakpoint-centered interval. For each chromatin dataset, overlap between TD-associated genomic intervals and ChIP-seq peaks was quantified and compared with a null distribution generated by genomic permutation. Chromatin association was expressed as the observed-to-expected overlap ratio, calculated as: observed overlap / mean overlap across randomized datasets (the same number of TDs per chromosome shuffled to random coordinates). An observed-to-expected ratio >1 indicates enrichment of TD-associated regions within the corresponding chromatin feature, whereas a ratio <1 indicates depletion. Empirical P values were calculated from the permutation distribution as the proportion of randomized datasets producing an overlap at least as extreme as the observed overlap in the tested direction.

### RNA-seq processing and expression quantification

RNA was isolated from baseline MDA-MB-436 cells and chronically FANCM-depleted cells across three independent biological replicates per condition using the RNeasy Mini Kit (Qiagen, cat. no. 74104). The resulting six mRNA-seq libraries, representing two experimental conditions with three biological replicates each, were sequenced as 100-bp single-end reads. RNA-seq data were processed using the Mayo Clinic MAP-RSeq pipeline v3.1.6. Reads were aligned to the GRCh38 human reference genome using STAR, and gene-level expression was quantified using Rsubread v2.1.1, generating both raw read counts and normalized expression values expressed as fragments per kilobase of transcript per million mapped reads (FPKM). Differential gene-expression analysis between baseline and chronically FANCM-depleted BRCA1-mutant MDA-MB-436 cells was performed using PyDESeq2, with raw gene-level counts as input. Differential expression was reported as |log2 fold change|>0.5 with significance assessed using multiple-testing-adjusted *P* values (false discovery rates, FDR). Fold changes were shrunk using DESeq2’s shrinkage method.

### Gene set enrichment analysis

Pathway-level transcriptional changes following chronic FANCM depletion were assessed by gene set enrichment analysis (GSEA) using a ranked list derived from the RNA-seq differential-expression results. Gene sets included Hallmark pathways and additional curated pathway collections represented in figure. Enrichment was summarized using the normalized enrichment score (NES), with positive NES values indicating enrichment among genes increased following FANCM depletion and negative NES values indicating enrichment among genes decreased following FANCM depletion. Multiple-testing correction was reported as the FDR q value. Pathways were visualized using bubble plots in which position represents NES, symbol size represents gene-set size and the indicated q value represents enrichment significance. Representative genes contributing to selected pathways were displayed in the accompanying expression heatmap.

### Integration of structural variants and gene expression

To determine whether newly emerged SVs were associated with local transcriptional changes, somatic SVs identified following chronic FANCM depletion were intersected with gene annotations and RNA-seq differential-expression results. Only genes meeting the final differential-expression criteria were classified as SV-associated differentially expressed genes.

SV–gene relationships were divided into three mutually interpretable spatial categories: Type A — gene fully contained within an SV. Genes whose complete annotated genomic span was encompassed by a deletion or duplication. Type B — SV breakpoint within the gene. Genes containing one or more breakpoints from a deletion, duplication or BND/rearrangement within the annotated gene body. Type C — breakpoint within 10 kb of the gene. Genes without an intragenic breakpoint but with an SV breakpoint located within 10 kb of the gene boundary, representing a proximal SV–gene relationship with potential regulatory consequences.

For each SV–gene relationship, expression direction among SV-associated genes was compared with a background consisting of genes with no identified relationship to the analyzed structural variants. Thus, the background rate represents transcriptional changes among SV-unaffected genes, rather than all genes in the transcriptome. This distinction was retained throughout the analysis to determine whether a given SV configuration was associated with expression direction beyond the genome-wide transcriptional response to chronic FANCM depletion.

Directional hypotheses were specified according to SV configuration: genes fully contained within deletions were tested for enrichment of decreased expression; genes fully contained within duplications were tested for enrichment of increased expression; genes directly disrupted by SV breakpoints were evaluated for decreased expression; genes located within 10 kb of SV breakpoints were evaluated for increased expression, where specified. For each SV configuration, the observed proportion of genes changing in the hypothesized direction was compared with the corresponding proportion among SV-unaffected background genes using a one-sided binomial test. The null probability for each test was therefore the observed directional-expression rate among SV-unaffected genes. Nominal P values were reported for these targeted comparisons.

### BRCA1-mutated breast cancer cohort selection, FANCM expression stratification and Structural variant calling

Breast cancer samples with somatic BRCA1 driver mutations or germline BRCA1 mutations were identified^1^. Samples with somatic driver or germline BRCA2 mutations were excluded. Additional breast cancer samples with BRCA1 single-nucleotide variants (SNVs), multi-nucleotide variants (MNVs), and small insertions and deletions (indels) were identified from the PCAWG final consensus somatic mutation call set^38^ (final_consensus_passonly.snv_mnv_indel.icgc.public.maf.gz), downloaded from the ICGC open-access data repository (s3://icgc25k-open/PCAWG/consensus_snv_indel/). However, these additional samples contained only silent, intronic, or intergenic BRCA1 variants and were not included. The remaining samples were matched to BRCA-EU samples with simple structural variant signature data^31^. One sample (PD14442a) was excluded because only two simple SVs were present in the SV signature data.

Breast cancer RNA-seq data generated as part of the BASIS consortium were provided by Dr. Marcel Smid^39^. The data were GeTMM-normalized^40^ and provided on a log2 scale. RNA and DNA samples from the same patient were matched using corresponding PR and PD identifiers. The final cohort included 15 BRCA1-altered breast cancer samples (13 germline and 2 somatic). FANCM expression was extracted and samples were stratified into FANCM-high (n=7) and FANCM-low (n=8) groups using a median split.

Structural variant calls for the 15 breast cancer WGS samples were obtained from the COSMIC bulk Structural Variants export. Each event was classified as DEL, DUP, INV or BND according to breakpoint strand orientation. The available COSMIC dataset contained structural variants >1 kb in size. SVs were pooled separately for FANCM-high and FANCM-low tumors and processed using the same size- and class-based framework applied to the experimental WGS data. Both absolute pooled event counts and proportional size distributions were visualized.

### Statistical Analysis

Statistical analyses were selected according to the experimental design and data type. SV architectural concordance between independently derived clones was assessed using Spearman’s rank correlation, Pearson’s correlation and R^2^, together with cosine similarity and Jensen–Shannon divergence for class-specific distributions.

Chromatin enrichment at TD breakpoint-associated regions was assessed by permutation testing against randomized genomic intervals. Differential RNA expression was analyzed using PyDESeq2 with FDR correction for genome-wide multiple testing. GSEA significance was assessed using FDR q values. Directional SV–gene expression associations were evaluated using one-sided binomial tests in which the null probability was defined by the corresponding expression-direction rate among SV-unaffected genes. For genome-wide analyses, adjusted P values or FDR q values were used as specified. For targeted SV–expression analyses, nominal P values are shown

All Statistical tests were performed on GraphPad Prism (version 10.6.1). Comparisons between two cell lines were conducted using unpaired Student’s *t*-test, and between more BRCA1-mutant and BRCA1-Wild-type groups (with at least 3 cell lines per group) using Nested-test. For analyses involving more than two groups, statistical significance was evaluated using one-way analysis of variance (ANOVA), followed by Tukey’s post hoc test for multiple pairwise comparisons. A p-value < 0.05 was considered statistically significant. Specific statistical tests and significance are provided in the corresponding figure legends.

## Acknowledgements

We thank Dr. Nagarajan Kannan and Syed Mohammed Musheer Aalam from Mayo Clinic for their assistance with the three-dimensional (3D) TNBC spheroid experiments. We thank Dr. Neil Johnson (Fox Chase Cancer Center, Philadelphia, PA) for providing the MDA-MB-436 *BRCA1* add-back cell line; Dr. John Hawse (Mayo Clinic, Rochester, MN) for providing the HCC1937 and Hs578T cell lines; and Dr. Daniel Durocher (University of Toronto, Toronto, Canada) for providing the RPE1-hTERT TP53^−/−^ and RPE1-hTERT TP53^−/−^ *BRCA1^−/−^*cell lines. We thank Dr. Marcel Smid from Erasmus University Medical Center, the Netherlands, for sharing RNA-seq data from the Breast Cancer Somatic Genetics Study (BASIS) consortium. We thank Dr. Krishna R. Kalari and Dr. Kevin Thompson from Mayo Clinic for their assistance with HRD score analysis. We thank Dr. Michael T. Lewis and the Patient-Derived Xenograft Core for characterizing the TNBC patient-derived xenograft (PDX) models.

This work was supported by the National Cancer Institute of the National Institutes of Health under award R00CA252044 to Arvind Panday, the Career Enhancement Program of the Mayo Clinic Breast Cancer SPORE (P50CA116201) to Arvind Panday, and a Pilot Project supported through the Mayo Clinic Comprehensive Cancer Center (MCCCC) under the National Cancer Institute Cancer Center Support Grant (P30CA015083) to Arvind Panday. The content is solely the responsibility of the authors and does not necessarily represent the official views of the National Institutes of Health.

## Author Contribution

F.R., U.S., S.B.S., A.V., N.Y., V.P., S.D., A.P. and S.L. conducted the experiments; F.R, U.S, S.B.S, A.P, H.V, S.K and A.P designed the experiments; F.R., Z.C., S.S. and A.P. analyzed the data; and F.R. and A.P. wrote the paper.

## Materials & Correspondence

All correspondence should be addressed to A.P.

---

**Supplementary Figure 1. Fraction of genome altered stratified by *BRCA1* status in breast cancer across independent cohorts.** FGA in *BRCA1*-mutant and *BRCA1*-wild-type breast tumors in invasive breast carcinoma cohorts TCGA *PanCancer Atlas* (n = 1050) **(a),** and TCGA *Nature 2012* (n = 482) **(b).** Asterisks denote significant p-value using two-tailed unpaired Student t-test (***p<0.001, *p<0.05).

**Supplementary Figure 2. Validation of FANCM knockdown and doxycycline-inducible FANCM depletion. a**, FANCM mRNA expression following FANCM knockdown in HCC1937, HCC1395, parental MDA-MB-436 and *BRCA1*-complemented MDA-MB-436 cells, measured by RT-qPCR. **b**, FANCM expression in inducible-shScrambled and shFANCM MDA-MB-436 cells cultured with or without DOX (2 µg /ml). Expression was normalized to GAPDH using the 2−ΔCT method. Asterisks denote significant p-value using two-tailed unpaired Student t-test (***p<0.001, *p<0.05, ns= non-significant).

**Supplementary Figure 3. Detailed span-size distributions of deletions and tandem duplications following FANCM depletion. a**, Baseline deletion counts across genomic span-size bins and grouped percentage distribution of deletions measuring 0-10 kb, 10-100 kb, 100 kb-1 Mb and >1 Mb. **b**, Corresponding size distributions of deletions newly emerging in shFANCM-Clone 1 and Clone 2. **c**, Baseline TD counts across genomic span-size bins and grouped percentage distribution of TDs measuring 0-10 kb, 10-100 kb, 100 kb-1 Mb and >1 Mb. **d**, Corresponding size distributions of TDs newly emerging in Clone 1 and Clone 2.

**Supplementary Figure 4. Concordance of structural variant architectures between independent FANCM-depleted clones. a**, Comparison of SV counts between shFANCM-Clone 1 and Clone 2 across class- and size-defined SV categories. Populated categories are displayed on logarithmic axes and colored by SV class. The dashed diagonal denotes equal counts between clones. Spearman’s ρ, Pearson’s *r* and R^2^ quantify concordance between the independently derived clones. **b**, Class-specific comparison of SV distributions between Clone 1 and Clone 2 using cosine similarity and Jensen-Shannon divergence (JSD). Higher cosine similarity and lower JSD indicate greater similarity between clones.

