## Supplement data for "FANCM restrains structural genome evolution and defines a synthetic lethal dependency in *BRCA1*-deficient breast cancer"

#### Supplement Figure 1

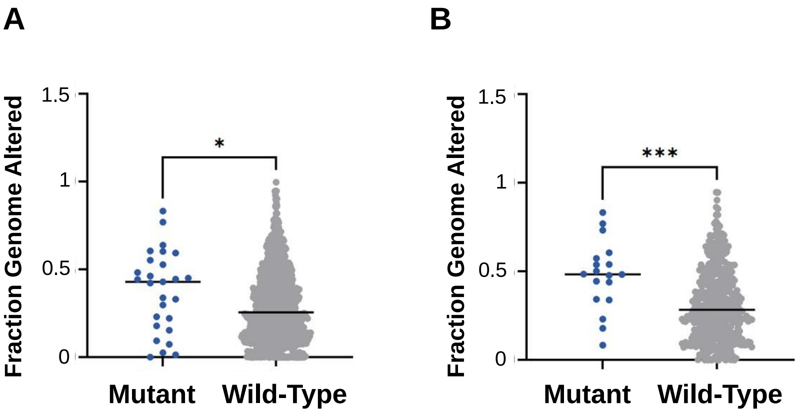

#### Supplement Figure 2

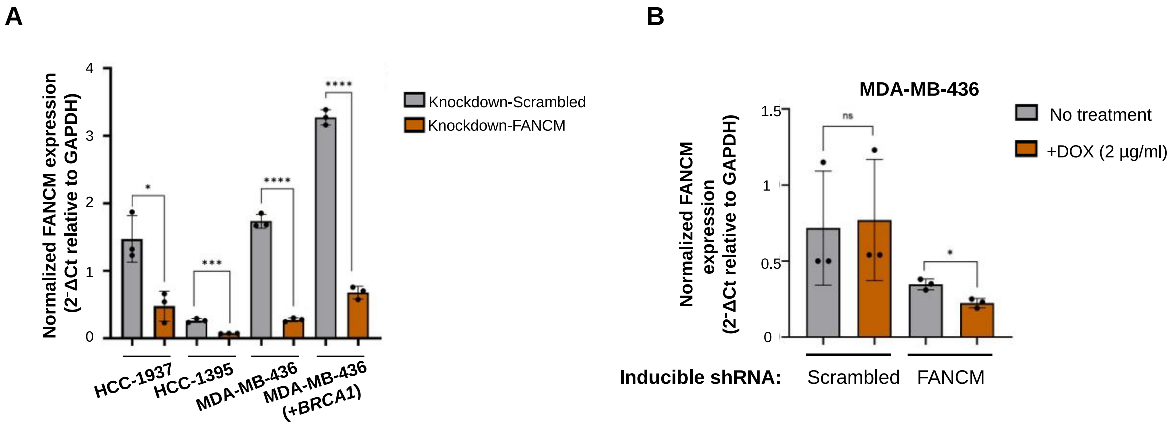

### Supplement Figure 3

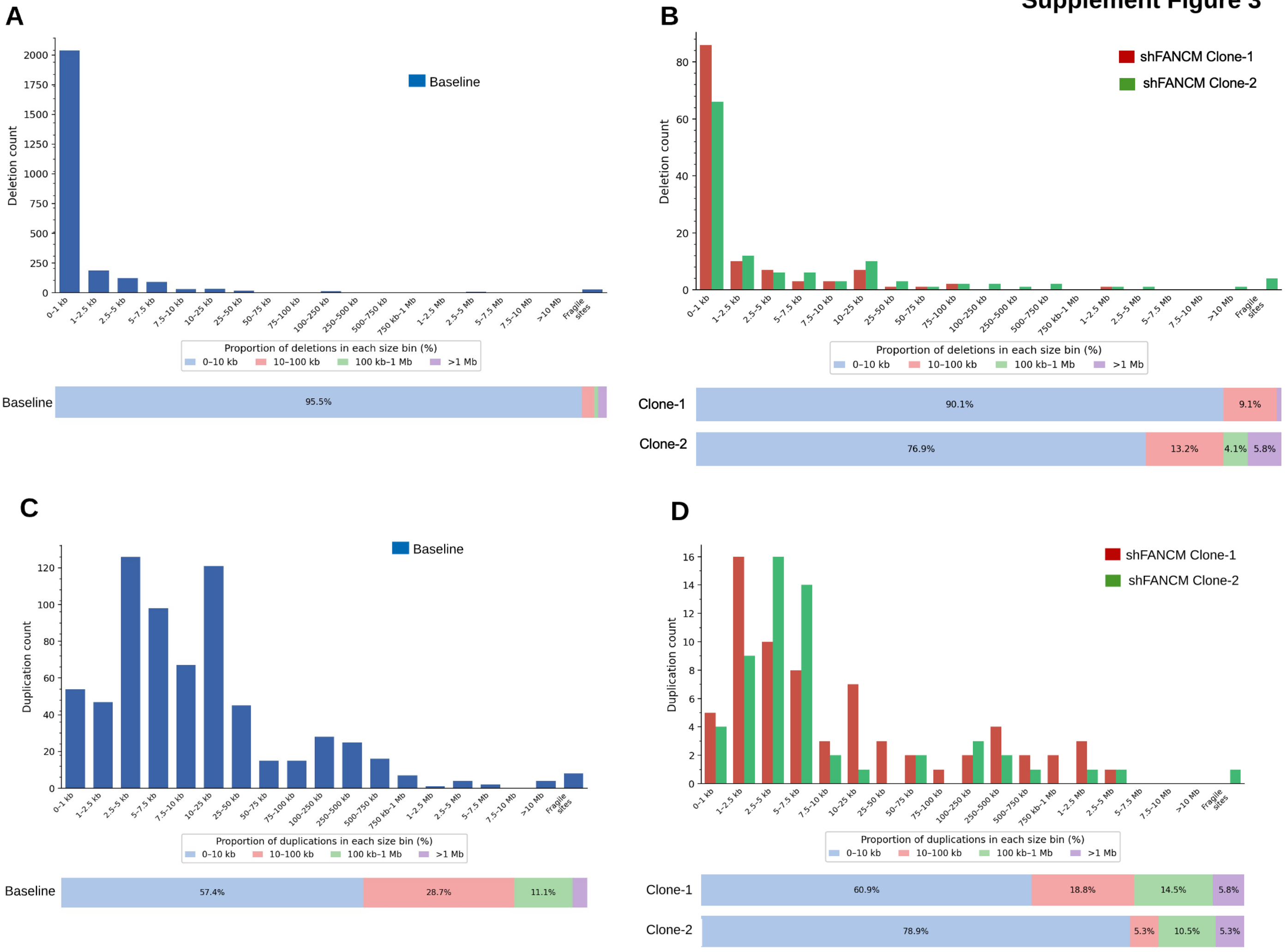

Supplement Figure 4

**A**

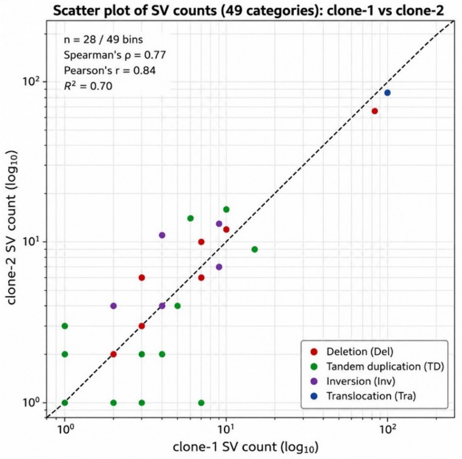


**B**

| Structural variant Type | Cosine similarity | Jensen-Shannon divergence (JSD) |
|---|---|---|
| **Deletion** | 0.9914 | 0.0421 |
| **Tandem Duplication** | 0.8373 | 0.0862 |
| **Inversion** | 0.8157 | 0.1074 |
| **Translocation** | 0.9987 | 0.003 |
